# Marked Synergy by Vertical Inhibition of EGFR signaling in NSCLC Spheroids: SOS1 as a therapeutic target in EGFR-mutated cancer

**DOI:** 10.1101/2020.04.28.066241

**Authors:** Patricia L. Theard, Erin Sheffels, Nancy E. Sealover, Amanda J. Linke, David J. Pratico, Robert L. Kortum

## Abstract

Drug treatment of 3D cancer spheroids more accurately reflects in vivo therapeutic responses compared to adherent culture studies. In EGFR-mutated lung adenocarcinoma, EGFR-TKIs show enhanced efficacy in spheroid cultures. Simultaneous inhibition of multiple parallel RTKs further enhances EGFR-TKI effectiveness. We show that the common RTK signaling intermediate SOS1 was required for 3D spheroid growth of EGFR-mutated NSCLC cells. Using two distinct measures of pharmacologic synergy, we demonstrated that SOS1 inhibition strongly synergized with EGFR-TKI treatment only in 3D spheroid cultures. Combined EGFR- and SOS1-inhibition markedly inhibited Raf/MEK/ERK and PI3K/AKT signaling. Finally, broad assessment of the pharmacologic landscape of drug-drug interactions downstream of mutated EGFR revealed synergy when combining an EGFR-TKI with inhibitors of proximal signaling intermediates SOS1 and SHP2, but not inhibitors of downstream RAS effector pathways. These data indicate that vertical inhibition of proximal EGFR signaling should be pursued as a potential therapy to treat EGFR-mutated tumors.

## Introduction

Lung cancer is the leading cause of cancer-related death worldwide; adenocarcinomas are the most common subtype of lung cancer. Oncogenic driver mutations in the RTK/RAS pathway are found in over 75% of lung adenocarcinomas (*1*). Activating EGFR mutations occur in 10-30% of lung adenocarcinomas, and are the major cause of lung cancer in never-smokers. In patients whose tumors harbor either an L858R mutation or an exon 19 deletion (85% of EGFR mutated tumors), first-generation EGFR-tyrosine kinase inhibitors (TKIs) erlotinib and gefitinib enhance progression free survival (*2–4*). However, resistance to first generation TKI invariably occurs. In most cases, acquired resistance to first generation EGFR-TKIs occurs via either a secondary EGFR ‘gatekeeper mutation’ (T790M, 50-60% of cases) that renders the receptor insensitive to first generation EGFR-TKIs or oncogenic shift to alternative RTKs (15-30%). To treat patients with T790M-mutated resistant tumors, the third generation EGFR-TKI osimertinib, which selectively targets activating EGFR mutant proteins including T790M but spares wild-type EGFR, was developed (*5, 6*). However, despite further enhancing survival of patients with EGFR-mutant tumors, resistance again emerges.

Unlike first-generation EGFR-TKIs, mechanisms driving osimertinib resistance are more variable, including both EGFR-dependent (10-30%) and EGFR-independent mechanisms (*7–10*). The most common EGFR-independent resistance mechanisms involve reactivation of the RTK/RAS/effector pathway (*10*), often via enhanced signaling through parallel RTKs (*7–9, 11–16*). Here, combining osimertinib with individual RTK inhibitors can both inhibit the development of resistance through the inhibited RTK and kill cancer cells with resistance driven by the specific RTK being inhibited. However, simultaneous inhibition of multiple RTKs with osimertinib may be required to eliminate oncogenic shift to alternative RTKs (*8*). Downstream of RAS, co-targeting intermediates of the RAF/MEK/ERK and PI3K/AKT pathways enhances of osimertinib effectiveness, however, signaling through the uninhibited effector pathway may drive resistance (*17–20*). Thus, it may be important for therapeutic combinations including osimertinib to stifle all downstream RTK/RAS signaling to be effective.

Recent studies suggest that pharmacologic assessments of targeted therapeutics should be performed under 3D culture conditions rather than in 2D adherent cultures (*21, 22*). 3D spheroids show altered growth characteristics, changes in cell surface proteins, altered metabolism, changes in activation of signaling pathways or altered responses to targeted pathway inhibitors, and are more resistant to drug-induced apoptosis compared to 2D adherent cultures signaling (*23–26*). These differences may be particularly relevant in *EGFR*-mutated NSCLC. *EGFR*-mutated cells show differential RTK expression and phosphorylation in 3D versus 2D conditions (*27*). Further, EGFR-mutated cells respond more robustly to first-generation EGFR-TKIs in 3D cultures, and these responses more closely resemble responses seen *in vivo* (*28*). These data highlight the need for pharmacologic assessment of therapeutics designed to treat *EGFR*-mutated NSCLC under 3D culture conditions.

The ubiquitously expressed RasGEFs (guanine nucleotide exchange factors) SOS1 and SOS2 (son of sevenless 1 and 2) are common signaling intermediates of RTK-mediated RAS activation. Although not initially considered as drug targets because of the low oncogenic potential of SOS (*29*), there has been renewed interest in SOS proteins as therapeutic targets for cancer treatment. We and others have shown that SOS1 and SOS2 may be important therapeutic targets in KRAS-mutated cancer cells (*30–32*), and a specific SOS1 inhibitor (BAY-293) has recently been identified (*33*). Here, we investigate SOS1 and SOS2 as potential therapeutic targets in EGFR-mutated lung adenocarcinoma cells. Using two distinct measures of pharmacologic synergy, we demonstrate that SOS1 inhibition using BAY-293 synergizes with osimertinib only under 3D spheroid culture conditions, and in doing so add to the growing evidence that pharmacologic assessment of novel therapeutics designed to treat cancer must be performed under 3D culture conditions (*27, 31, 34–36*). By assessing the pharmacologic landscape of EGFR/RAS pathway inhibitors, we demonstrate that inhibition of proximal signaling is required to synergize with osimertinib, and that combined EGFR and SOS1 inhibition synergizes to inhibit RAS effector signaling in 3D culture. These findings have significant therapeutic implications for the development of combination therapies to treat EGFR-mutated lung adenocarcinoma.

## Results

### SOS1 deletion inhibits transformation in EGFR-mutated NSCLC cells

Previous studies showed that EGFR-mutated NSCLC cell lines show much more robust responsiveness to first generation EGFR-TKIs in 3D (spheroid or organoid) culture compared to 2D adherent culture, and further that 3D conditions more readily mirror EGFR-TKI responses seen *in vivo* (*28*). To confirm these findings and extend them to third generation EGFR-TKIs, we assessed dose-dependent survival of both first-generation EGFR-TKI sensitive (HCC827, exon 19 deletion [Δ*ex*19]) or resistant (NCI-H1975, L858R/T790M) NSCLC cell lines to either gefitinib or osimertinib treatment under both adherent (2D) or spheroid (3D) culture conditions (Fig. 1A). HCC827 and H1975 cells were plated in either adherent or spheroid cultures, allowed to rest for 48 hours, and then treated with increasing doses of either the first-generation EGFR-TKI gefitinib or the third-generation EGFR-TKI osimertinib for four days. HCC827 cells showed responsiveness to both EGFR-TKIs under 2D and 3D culture conditions, however in both cases 3D spheroid cultures showed a > 1-log enhancement in EGFR-TKI efficacy and enhanced overall growth inhibition. While NCI-H1975 cells were not sensitive to gefitinib, osimertinib treatment of H1975 cells showed enhanced efficacy and increased overall growth inhibition in 3D spheroids over 2D adherent cultures.

**Figure 1.** *SOS1* deletion inhibits anchorage-dependent (3D) transformation in EGFR-mutated NSCLC cell lines. **(A)** Dose-response curves of EGFR-mutated HCC827 (Δex19) (left) or NCI-H1975 (L858R/T790M) (right) cells treated with gefitinib or osimertinib under 2D anchorage-dependent (gray diamonds) or 3D spheroid (black squares) culture conditions. **(B-C)**2D proliferation (left) or 3D spheroid growth (right) in **(B)** HCC827 or **(C)** NCI-H1975 cells where *SOS1* or *SOS2* has been deleted using CRISPR/Cas9 vs NT controls. 10x images of representative spheroids at day 0 and 21 are shown, scale bar=250 mm. **(D)** 3D transformation in the indicated EGFR-mutated NSCLC cell lines where *SOS1* or *SOS2* has been deleted using CRISPR/Cas9 vs NT controls. **(E)** Dose-response curve cells of NCI-H1975 cells treated with the SOS1 inhibitor BAY-293 under 2D anchorage-dependent (gray diamonds) or 3D spheroid (black squares) culture conditions. **(F)** Dose-response curves of NCI-H1975 cells where *SOS1* (red circles) or *SOS2* (blue triangles) has been deleted using CRISPR/Cas9 vs NT controls (black squares) treated with BAY-293 under 3D spheroid culture conditions. Dose-response curves and 2D proliferation are presented as mean +/− s.d. from three independent experiments. For transformation studies, data are from four independent experiments. Statistical significance was determined by ANOVA using Tukey’s method for multiple comparisons. * p<0.05, **p<0.01, ***p<0.001 vs. NT cells. # p<0.05, ##p<0.01 vs. *SOS1* KO cells.

SOS1 and SOS2 are ubiquitously expressed RasGEFs responsible for transmitting EGFR signaling to downstream effector pathways. To determine whether SOS1 or SOS2 were required for 2D anchorage-dependent proliferation or 3D spheroid growth in EGFR-mutated NSCLC cells, *SOS1* ((*37*) and Fig. S1) or *SOS2* (*31*) were deleted in pooled HCC827 and H1975 cells, and both proliferation and spheroid growth were assessed versus NT controls (Fig. 1B and C). In adherent culture, neither *SOS1* nor *SOS2* deletion altered proliferation (Fig. 1B). In contrast, *SOS1* deletion completely inhibited spheroid growth in both HCC827 and H1975 cells, indicating that SOS1 was required to maintain the transformed phenotype in both cell lines. To determine whether SOS1 was generally required for mutant EGFR-driven transformation, we further deleted *SOS1* or *SOS2* in both first-generation sensitive NCI-H3255 (L858R) and PC9 (Δ*ex*19) cells and in subcultures of these cell lines that had acquired T790M mutations after continuous EGFR-TKI treatment (H3255-TM (*38*) and PC9-TM (*39*)). In all cases, *SOS1* deletion significantly diminished oncogenic transformation, whereas *SOS2* deletion had variable effects on transformation depending on the EGFR mutated cell line examined (Fig. 1D). These data indicate that SOS1 is the major RasGEF responsible for oncogenesis downstream of mutated EGFR.

BAY-293 was recently described as a specific inhibitor for SOS1 (*33*). To determine whether SOS1 inhibition was similarly more effective in 3D spheroids over 2D adherent culture, we assessed dose-dependent survival of H1975 cells after BAY-293 treatment under both 2D and 3D culture conditions (Fig. 1E). Similar to what we observed after either EGFR-TKI treatment (Fig. 1A) or *SOS1* deletion (Fig. 1C and D), BAY-293 showed enhanced efficacy and increased overall growth inhibition in 3D spheroids over 2D adherent cultures. To confirm the specificity of BAY-293 for SOS1, we further treated 3D spheroid cultured H1975, PC9-TM, and H3255-TM cells where either *SOS1* or *SOS2* had been deleted versus NT controls with increasing doses of BAY-293 (Fig. 1F and Fig. S2). BAY-293 treatment did not inhibit survival of spheroids where *SOS1* had been deleted, indicating the specificity of BAY-293 for SOS1. Further, cells where *SOS2* had been deleted showed an approximately 1-log enhancement in BAY-293 efficacy and enhanced overall growth inhibition compared to NT controls, indicating that SOS1 and SOS2 have some overlapping functions in supporting survival of spheroid cultured EGFR-mutated NSCLC cells. Overall, these data suggest that EGFR-mutated NSCLC cells are more sensitive to either mutant EGFR or SOS1 inhibition in 3D spheroid culture compared to traditional 2D adherent conditions.

### SOS1 inhibition synergizes with EGFR-TKIs to inhibit cell survival under anchorage independent (3D) culture conditions

Previous studies reported that combining osimertinib with an alternative RTK inhibitor may inhibit or treat the development of resistance driven by that specific RTK (*7–9*), whereas simultaneous inhibition of multiple parallel RTKs with osimertinib may be required to effectively potentiate osimertinib action (*8*). Further, while many studies show enhanced drug activity in combination therapies versus osimertinib treatment alone, they do not assess whether the effects of the 2-drug combinations are truly synergistic; synergistic interactions between therapeutics allow for maximization of the therapeutic effect while minimizing adverse events and may be required for effective therapeutic combinations with targeted agents (*40*).

SOS1 is a common downstream mediator of RTK signaling. We hypothesized that SOS1 could be an effective drug target to synergize with EGFR-TKI inhibition to treat EGFR-mutated lung adenocarcinoma. To directly assess synergy between osimertinib and SOS1 inhibition, we use two distinct methods based on the most widely established reference models of drug additivity. The first method, isobologram analysis, assesses changes in the dose-response curves for mixtures of two drugs compared to sham mixtures of each individual drug with itself. The second method, Bliss independence analysis, assesses whether a mixture of two individual drug doses has a greater effect than would be expected if the two drugs acted independently. We will first describe and then use each method in turn to determine the whether SOS1 inhibition using BAY-293 could synergize with the EGFR-TKI osimertinib in *EGFR*-mutated lung adenocarcinoma cells.

Isobologram analysis is a dose-effect analysis based on the principle of Loewe additivity, which states that a drug mixed with itself, and by extension a mixture of two or more similar drugs, will show additive effects. For two drugs (Drug A and Drug B) that have parallel dose-response curves so that a constant potency ratio is maintained at all doses of A and B (Fig. 2A), treatment using any dose-equivalent (DEQ) mixture of Drugs A and B will show a similar effect to treatment with either Drug A or Drug B alone if the effects of the two drugs are additive. In contrast, if the two drugs show synergism, then the effect seen by treatment with DEQ mixtures of A and B will be greater than the effect for either drug alone. By generating dose-response curves for different DEQ mixtures of Drugs A and B (Fig. 2B), one can compare the EC_50_ of each DEQ mixture to the EC_50_ of Drug A or Drug B alone on an isobologram plot (Fig. 2C). The EC_50_ of each individual drug is plotted as the x- or y-intercept, and the calculated contribution of each drug to the overall EC_50_ for each DEQ mix is plotted as a single point (EC_50,A_, EC_50,B_) on the graph. If the EC_50_ values for each DEQ mix fall along the straight line (isobole) that connects the individual drug EC_50_ values, then the drug-drug interaction is additive. In contrast, points that fall above or below the isobole indicate antagonism or synergy. The extent to which two drugs interact can be further quantified from the EC_50_ data as a combination index (CI) (Fig. 2D). A CI between 0.8-1.2 indicates the two drugs have additive effects when combined, a CI < 0.8 indicates synergy, and a CI > 1.2 indicates antagonism.

**Figure 2.**
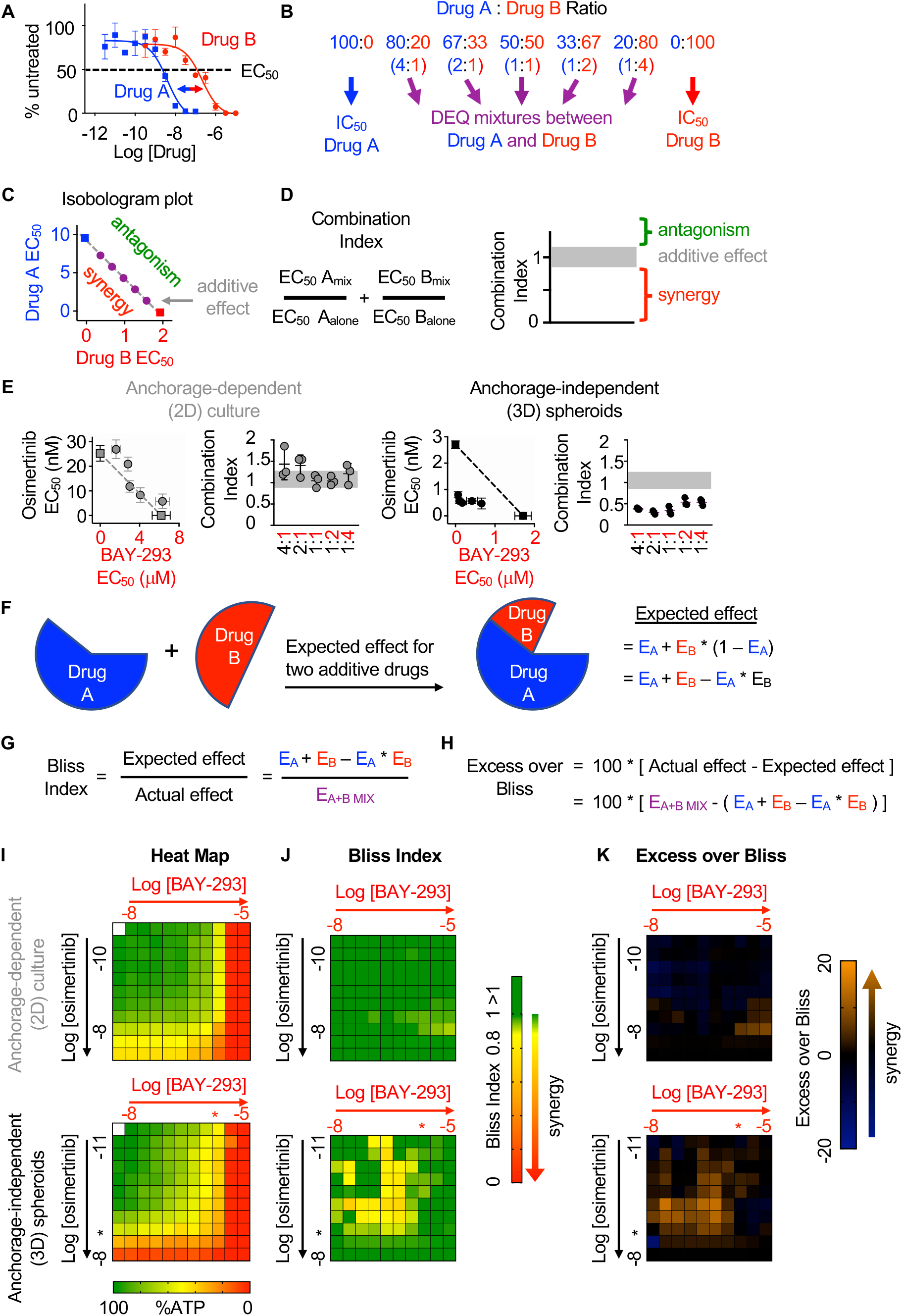
SOS1 inhibition synergizes with the EGFR-TKI inhibitor osimertinib to inhibit cell survival under anchorage-independent (3D) culture conditions. **(A-D)** Isobologram analysis examines drug-drug synergy by comparing dose equivalent (DEQ) mixtures of two drugs based on their EC_50_ values to treatment with either drug alone **(A and B)**. From the dose-response curves of the DEQ mixtures, plotting the fractional EC_50_ for each drug in the combination (purple) relative to the individual drug EC_50_ values (blue, red) on an isobologram plot **(C)** and calculation of the combination index (CI, **D and E**) allows assessment of drug-drug synergy. Additive effects occur on the dashed lines of the isobologram plot and have a CI 0.8-1.2 (gray box), whereas synergistic interactions fall below the dashed lines and have a CI < 0.8. **(E)** Isobologram plots and CI from dose-equivalent treatments of H1975 EGFR-mutated NSCLC cells treated with DEQ combinations of osimertinib and BAY-293. Isobologram and CI data are presented as mean +/− s.d. from three independent experiments. **(F)** Bliss additivity evaluates whether the overall effect of an individual drug combination (E_A+B_ mix) is greater than should be expected for two drugs with independent effects on the overall population (E_A_ + E_B_ – E_A_ * E_B_). **(G)** The Bliss Index compares the ratio of the expected effect to the actual effect. Synergistic interactions have a Bliss Index < 0.85. **(H)** Excess over Bliss evaluates the magnitude of the difference between the actual and expected effects. Increasingly synergistic interactions show an excess over Bliss Index > 0. **(I)** Heat map of H1975 cells treated with the indicated doses of osimertinib and/or BAY-293 grown in either 2D (adherent) culture conditions or as 3D spheroids. Green indicates more cells, red indicates fewer cells. EC_50_ values for each individual drug are indicated by an *. **(J)** Heat map of Bliss Index assessing drug-drug synergy between osimertinib and BAY-293 at each dose combination from **D**. **(K)** Heat map of excess over Bliss assessing drug-drug synergy between osimertinib and BAY-293 at each dose combination from **D**. Bliss Index and excess-over Bliss are presented as the mean from three independent experiments.

To assess drug-drug synergy between osimertinib and BAY-293 via isobologram analysis, NCI-H1975 cells were cultured under 2D adherent or 3D spheroid conditions for 48 hours, and were treated with varying DEQ combinations of osimertinib:BAY-293 (see Fig. 2B) for four days. Cell viability data was assessed using CellTiter-Glo and EC_50_ values from each DEQ mixture were used to generate isobologram plots and calculate combination indices (Fig. 2E). When cells were cultured under 2D conditions, osimertinib and BAY-293 showed additive effects, as DEQ EC_50_ values fell on the isobole and CI values were between 0.8-1.2. In contrast, when cells were cultured as 3D spheroids, osimertinib and BAY-293 showed significant synergy, as DEQ EC_50_ values were well below the isobole and CI < 0.8.

Bliss independence analysis is an effect-based analysis based on the principle of Bliss additivity, which assumes that two drugs will act independently of each other so that their combined effect can be assessed by assessing the effect of each drug sequentially (Fig. 2F). Unlike isobologram analysis, this method does not require that two drugs being assessed have parallel dose-response curves and can be calculated based as few as three drug treatments, the effect each drug has on its own on the cell population, and the effect of combining the two drug treatments together. By representing the effect of each drug treatment as a probabilistic outcome between 0 (no effect) and 1 (100% effect), we can compare the observed effect of the drug-drug combination to the expected effect if each drug acted independently (Fig. 2E). The ratio of the expected effect to the observed effect is the Bliss Index (BI), where a BI < 1 indicates synergy (Fig. 2G). Alternatively, the magnitude of the difference between the observed and expected result can be reported as the excess over Bliss (Fig. 2H). While excess over Bliss is the most widely reported synergy metric, the Bliss Index can be directly compared with the combination index in isobologram experiments and should be used when both synergy methods are used to assess a given drug-drug interaction.

To assess drug-drug synergy between osimertinib and BAY-293 via Bliss Independence analysis, NCI-H1975 cells were cultured under 2D adherent or 3D spheroid conditions for 48 hours and were treated with increasing doses of BAY-293, osimertinib, or combinations of the two drugs over a 3-log scale for four days. Cell viability was determined using CellTiter-Glo and overall viability (Fig. 2I), Bliss index (Fig. 2J), and excess over Bliss (Fig. 2K) were represented as heat-maps. Similar to what we observed for isobologram analysis, osimertinib and BAY-293 did not show significant synergy in cells cultured under 2D adherent conditions. In contrast, we observed significant synergy between osimertinib and BAY-293, mostly at dose combinations of osimertinib and BAY-293 falling just below the individual drug EC_50_ values. Overall, the data presented in Fig. 2 indicate that osimertinib and BAY-293 show significant drug-drug synergy in EGFR-mutated H1975 cells, but only in 3D spheroid culture conditions.

To determine whether the SOS1 inhibitor BAY-293 could generally synergize with EGFR-TKIs in EGFR-mutated lung adenocarcinoma cells, we extended our assessment of drug-drug synergy to isobologram analysis (Fig. 3) and Bliss independence analysis (Fig. 4) in six different EGFR-mutated lung adenocarcinoma cell lines. In cells that were sensitive to first-generation EGFR-TKIs (HCC827, PC9, H3255; T790 wild-type), we assess drug-drug synergy between BAY-293 and either a first-generation (gefitinib) or third-generation (osimertinib) EGFR-TKI. In cells that were resistant to first-generation EGFR-TKIs (H1975; PC9-TM, H3255-TM; T790M) we limited our assessment to synergy between BAY-293 and osimertinib. To first determine the individual EC_50_ values for gefitinib, osimertinib, and BAY-293 in each cell line, cells were culture as 3D spheroids for 48-72 hours, and then treated with increasing doses of drug for four days followed by assessment of cell viability by CellTiter-Glo (Fig. S3). In five of six cell lines, the individual dose-response curves for BAY-293, osimertinib, and gefitinib (where appropriate) showed similar maximal effects and Hill coefficients, and were thus appropriate for linear isobologram analysis for each 2-drug combination of BAY-293, osimertinib, and gefitinib (*41*). In contrast, H3255-TM cells were only moderately sensitive to osimertinib, showing at most a 50% reduction in viability at high doses. Therefore, we limited our assessment of drug-drug synergy in H3255-TM cells to Bliss independence analysis. Further, to simplify our assessment of Bliss independence across multiple drugs and cell lines, we limited our drug treatments to 1:2, 1:1, and 2:1 mixtures of each drug combination based on dose equivalence (see Fig. 4A).

**Figure 3.**
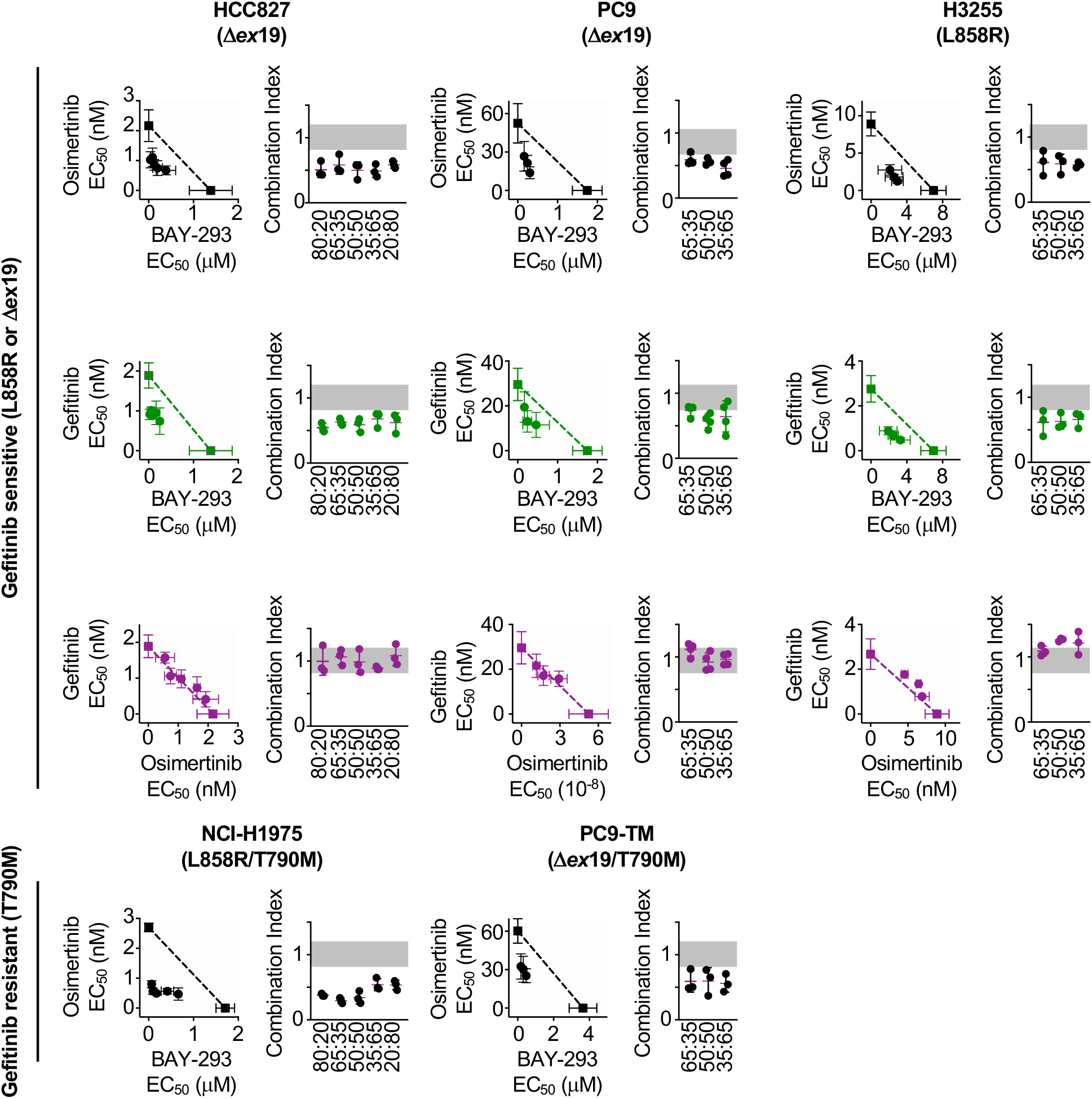
Isobologram analysis showing that SOS1 inhibition synergizes with EGFR-TKI treatment to inhibit survival in multiple EGFR-mutated NSCLC cell lines. Isobologram analysis and Combination Index (CI) from dose-equivalent treatments of the indicated EGFR-mutated gefitinib-sensitive (L858R or Dex19, top) or gefitinib-resistant (T790M, bottom) NSCLC cell lines with combinations of gefitinib, osimertinib, and BAY-293. Additive effects occur on the dashed lines of the isobologram plot and have a CI 0.8-1.2 (gray box), whereas synergistic interactions fall below the dashed lines and have a CI < 0.8. Data are presented as mean +/− s.d. from three independent experiments.

**Figure 4.**
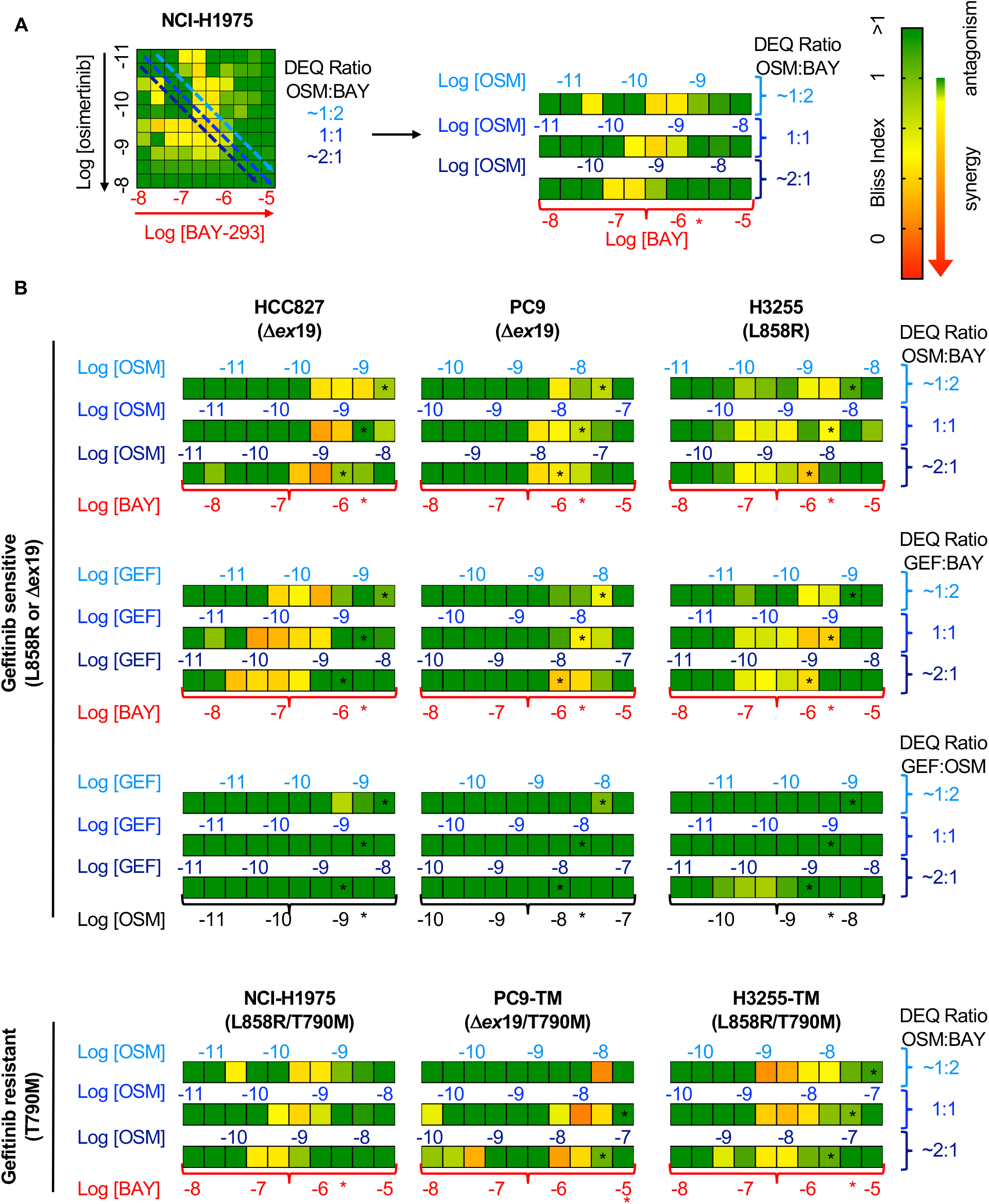
Bliss Independence analysis showing that SOS1 inhibition synergizes with EGFR-TKI treatment to inhibit survival in multiple EGFR-mutated NSCLC cell lines. **(A)** Bliss Index heatmap from 3D spheroid cultured NCI-H1975 cells Fig. 2A (left) and horizontal projections of Bliss Indices of drug treatments at 2:1, 1:1, and 1:2 ratios of osimertinib:BAY-293 based on dose equivalencies (right). Increasingly synergistic interactions (Bliss index < 0.85) are indicated by the corresponding heat map. The concentration of BAY-293 (held constant, bottom) and of osimertinib (above each horizontal projection) are given. The IC_50_ for each individual drug are shown (*). **(B)** Bliss Index heatmaps based on A for the indicated gefitinib-sensitive and gefitinib-resistant cell lines at 2:1, 1:1, and 1:2 ratios of osimertinib, gefitinib, and BAY-293 based on dose equivalencies. Data for NCI-H1975 cells are the same as in A. Data are presented as the mean from three independent experiments.

For each first-generation EGFR-TKI sensitive cell line (HCC827, PC9, H3255), gefitinib and osimertinib did not show any synergy with each other by either isobologram analysis (Fig. 3) or Bliss Independence analysis (Fig. 4), instead showing additive effects (CI and BI ~ 1) as would be expected for two drugs with the same molecular target. In contrast, BAY-293 showed significant synergy with gefitinib and osimertinib by both isobologram analysis (Fig. 3) and Bliss Independence analysis (Fig. 4), suggesting that SOS1 inhibition can act as a secondary treatment for all EGFR-TKIs. Further, in all three T790M mutated cell lines (H1975, PC9-TM, H3255-TM), BAY-293 again showed synergy with osimertinib. These data suggest that combined SOS1 and EGFR inhibition is a robust therapeutic combination that synergize to inhibit EGFR-mutated lung adenocarcinoma cell growth.

### Synergy between BAY-293 and osimertinib is independent of SOS2

We showed that *SOS2* deletion sensitized NCI-H1975 cells to the SOS1 inhibitor BAY-293 (Fig 1F). We wanted to determine whether the synergy we observed between EGFR- and SOS1-inhibition (Fig. 3 and 4) was enhanced by *SOS2* deletion in EGFR-mutated NSCLC cell lines. To examine whether SOS2 deletion alters the synergy between osimertinib and BAY-293 in EGFR (T790M) mutated cells, SOS2 was deleted in H1975, PC9-TM, and H3255-TM cells. For H1975 and PC9-TM cells, *SOS2* KO cells vs NT controls were cultured under 3D spheroid conditions for 48-72 hours, and were then treated with varying DEQ combinations of osimertinib:BAY-293 for four days. Cell viability data was assessed using CellTiter-Glo and EC_50_ values from each DEQ mixture were used to generate Isobologram plots and calculate confidence intervals (Fig. 5A and B). For both cell lines, *SOS2* deletion sensitized cells to BAY-293, decreasing EC_50_ by 5-10-fold compared to NT controls without altering the EC50 to osimertinib treatment alone. However, unlike what we observed in the NT control cells, osimertinib and BAY-293 showed only mild synergy in EGFR-mutated cells where *SOS2* was deleted as assessed by the distance of the interaction points to the isobole and the increased combination index vs. NT controls. Further, when we overlaid the NT and *SOS2* KO isobologram plots at two different scales of BAY-293, the drug combination data points were overlapping between NT and *SOS2* KO cells, suggesting that *SOS2* deletion did not enhance synergy between osimertinib and BAY-293.

**Figure 5.**
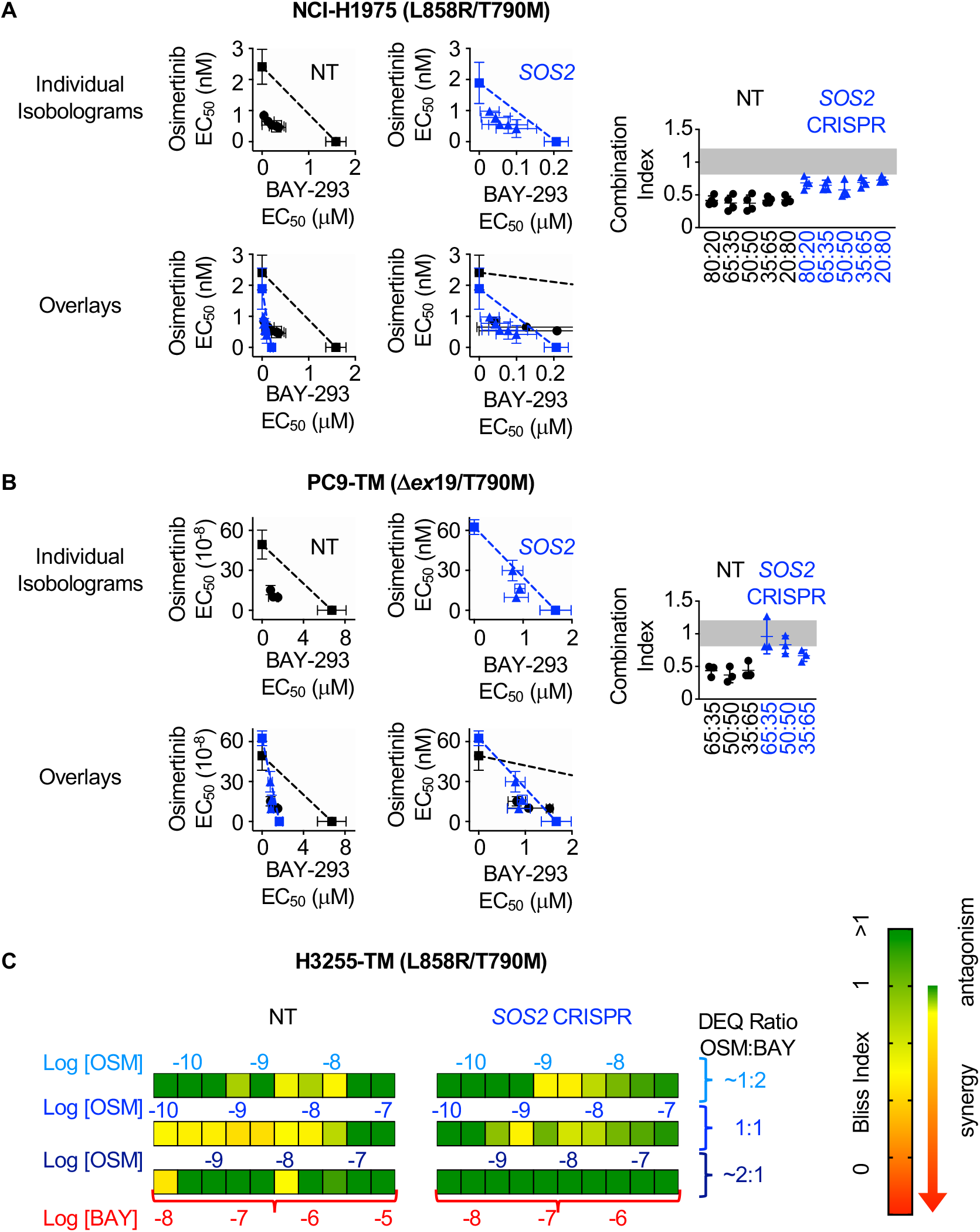
*SOS2* deletion does not enhance the synergistic interaction between SOS1 inhibition and EGFR-TKI treatment. **(A-B)** Isobologram analysis (left) and Combination Index (right) from dose-equivalent treatments of osimertinib and BAY-293 in H1975 (**A**) or PC9-TM (**B**) cells where *SOS2* has been deleted (blue) versus NT controls (black). Overlay plots on two different BAY-293 dosing scales are shown below the individual isobologram plots. Additive effects occur on the dashed lines of the isobologram plot and have a CI 0.8-1.2 (gray box), whereas synergistic interactions fall below the dashed lines and have a CI < 0.8. **(C)** Bliss Index heatmaps for H3255-TM cells where *SOS2* has been deleted versus NT controls treated at at 1:2, 1:1, and 2:1 ratios of osimertinib and BAY-293 based on dose equivalencies. Data are presented as mean +/− s.d. from three independent experiments.

Since H3255-TM cells are not appropriate for linear isobologram analysis between BAY-293 and osimertinib, we instead performed Bliss independence analysis to assess potential synergy between osimertinib and BAY-293 in the presence or absence of SOS2. H3255-TM cells where *SOS2* had been deleted vs NT controls were cultured under 3D spheroid conditions for 48-72 hours, and were then treated with increasing doses of osimertinib alone, BAY-293 alone, or mixtures of each drug dose at 1:2, 1:1, and 2:1 mixtures of osimertinib and BAY-293 based on dose equivalence for four days. Cell viability data was assessed using CellTiter-Glo, and the Bliss index was calculated for each drug mixture as shown in Fig. 2C and Fig. 4. As was the case in H1975 and PC9-TM cells, while the *SOS2* deletion sensitized H3255-TM cells to BAY-293 we observed less overall synergy between osimertinib and BAY-293 H3255-TM cells where we had deleted *SOS2* vs NT controls. These data suggest that although osimertinib and BAY-293 synergize to limit viability of EGFR-mutated lung adenocarcinoma cells, the synergy between osimertinib and BAY-293 is independent of SOS2.

### BAY-293 and osimertinib synergize to inhibit RAS effector signaling

Mutated EGFR signals through downstream RAF/MEK/ERK and PI3K/AKT effector pathways to promote proliferation, transformation, and survival. Since *SOS2* deletion did not further enhance synergy between BAY-293 and osimertinib, we hypothesized that SOS1 inhibition specifically enhanced EGFR-TKI dependent inhibition of downstream signaling in 3D culture. To perform signaling experiments on 3D cultured spheroids, cells were seeded in 24-well micropatterned low-attachment culture plates (Aggrewell, StemCell) containing ~1200 individual spheroids per condition. To determine the extent to which SOS1 inhibition and/or *SOS2* deletion altered osimertinib-dependent inhibition of downstream effector signaling in 3D culture, H1975 or PC9 cells where *SOS2* was deleted vs. NT controls were cultured as spheroids for 48-72 hrs and then treated with increasing doses of osimertinib +/− BAY-293 prior to spheroid collection, lysis, and Western blotting for phosphorylated ERK and AKT (Fig. 6). In both NT and *SOS2* knockout cells, BAY-293 reduced the dose of osimertinib required to inhibit both ERK and AKT phosphorylation (Fig. 6). For Raf/MEK/ERK signaling, Bliss Independence analysis of pERK quantitation revealed that either SOS1 inhibition or *SOS2* deletion independently synergized with osimertinib to inhibit Raf/MEK/ERK signaling, and the combination of inhibiting SOS1/2 signaling further enhanced this synergy. In contrast, for PI3K/AKT signaling *SOS2* deletion did not enhance the synergy between osimertinib and BAY-293. While either osimertinib treatment or *SOS2* deletion independently synergized with BAY-293 to inhibit AKT phosphorylation, *SOS2* deletion did not further enhance the ability osimertinib to inhibit PI3K/AKT signaling in the presence or absence of BAY-293. These data strongly suggest that vertical inhibition of EGFR and SOS1 limits call viability by inhibiting activation of both RAF/MEK/ERK and PI3K/AKT effector pathways.

**Figure 6.**
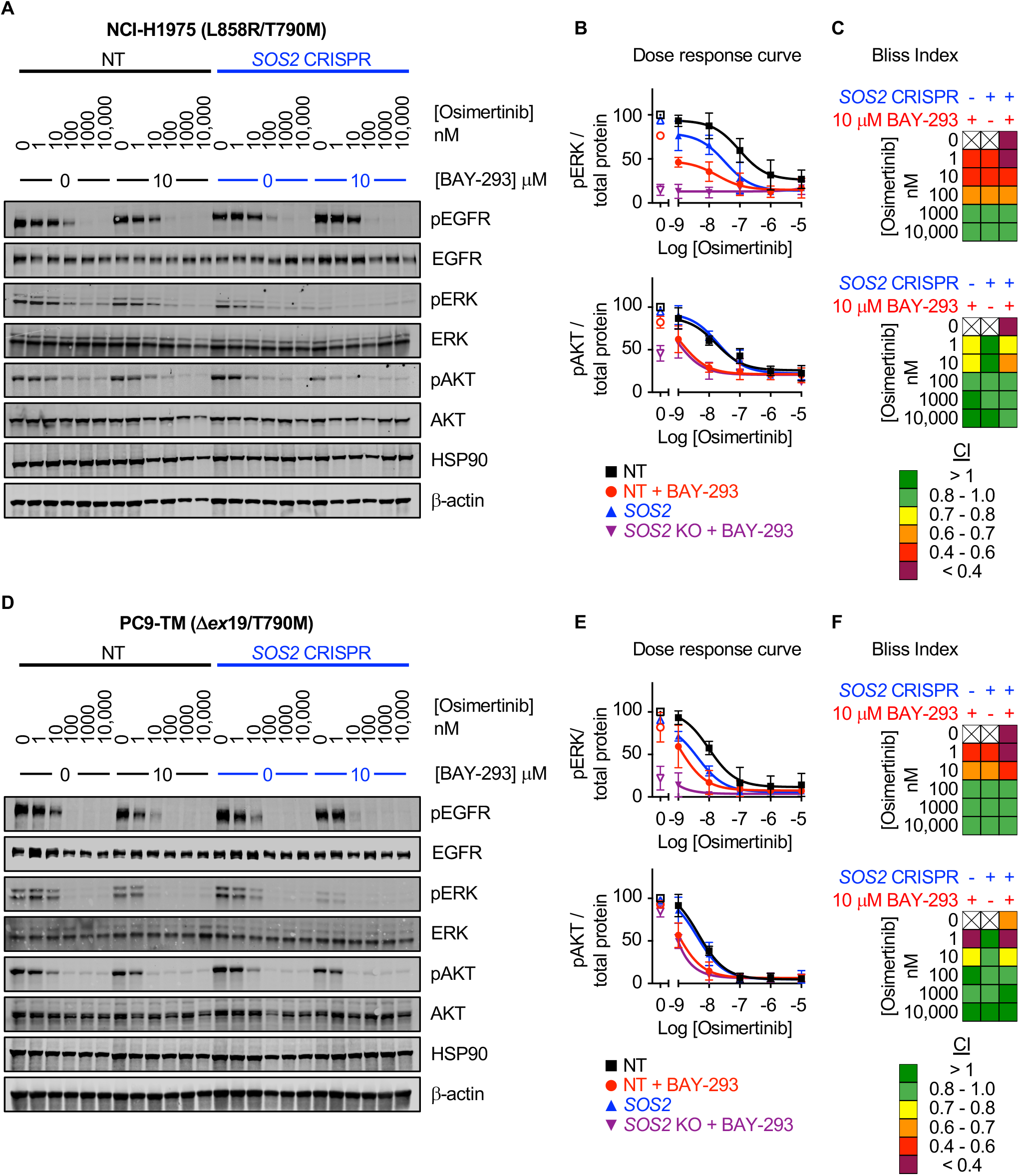
SOS1 inhibition synergizes with mutant EGFR inhibition to inhibit downstream effector signaling. Western blots **(A, D)**, pERK and pAKT quantitation **(B, E)**, and Bliss Indices **(C, F)** of WCLs of NCI-H1975 cells (A-C, top) or PC9-TM cells (D-F, bottom) cultured under 3D spheroid conditions for 48 hours and then treated with the indicated concentrations of the EGFR-TKI osimertinib and/or the SOS1 inhibitor BAY-293 for 6 hours. Western blots are for pEGFR, EGFR, pAKT, AKT, pERK1/2, ERK1/2, HSP90, and β-actin. pERK and pAKT quantifications were calculated using a weighted average of total protein western blots. Combination Indices are based on pERK / Total protein and pAKT / Total protein quantitations. Increasingly synergistic combinations are indicated in yellow, orange, red, or purple. Phosphoprotein quantitations are presented as mean +/− s.d. from three independent experiments. Bliss indices are presented as mean from three independent experiments.

### Assessment of Inhibitor Landscape in EGFR-mutated cells lines shows synergy upon inhibition of upstream pathway effectors

Since the most common EGFR-independent resistance mechanisms involve reactivation of RTK/RAS/effector pathways (*7–10*), we wanted to assess whether inhibition of different proteins within the EGFR/RAS signaling pathway could synergize to inhibit 3D survival of EGFR (T790M) mutated cancer cells. To determine drug-drug synergies after inhibition of EGFR-RAS pathway signaling at different levels, we assessed synergy between osimertinib, inhibitors of EGFR signaling intermediates of RAS (BAY-293 for SOS1 and RMC-4450 for SHP2), and inhibitors of the Raf/MEK/ERK (trametinib) and PI3K/AKT (buparlisib) pathways (Fig. 7A). H1975 and PC9-TM cells were treated with each individual inhibitor or 1:1 DEQ mixtures of every drug-drug combination, and the combination index was calculated to assess drug-drug synergy. Since 3255-TM cells are not suitable for isobologram analysis, these cells were treated with full-dose mixtures based on dose equivalence and the Bliss Index was calculated for each drug-drug combination (Fig. 7B). Intriguingly, all three cell lines showed drug-drug synergy with any combination of EGFR, SOS1, and SHP2 inhibition. In contrast, inhibition of downstream Raf/MEK/ERK or PI3K/AKT pathways failed to consistently synergize with either osimertinib or any other inhibitor (Fig. 7B, top). These data support the premise that combined vertical inhibition of proximal EGFR signaling may constitute an effective strategy to treat EGFR-mutated lung adenocarcinomas.

**Figure 7.**
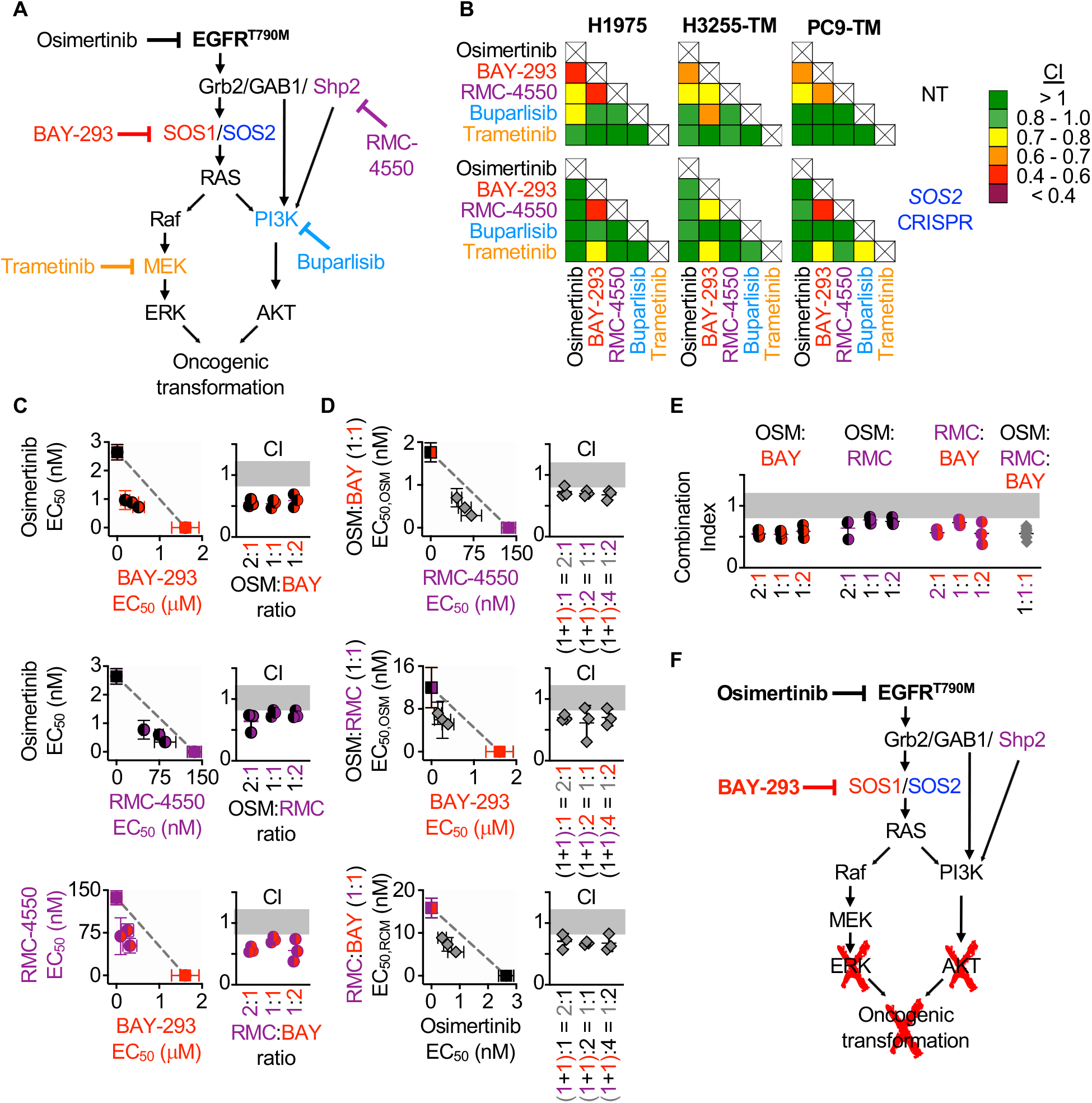
Assessment of the EGFR/RAS pathway ‘inhibitor landscape’ suggests that combination therapies inhibiting mutated EGFR, SOS1, and SHP2 have therapeutic potential in EGFR-mutated NSCLC. **(A)** Signaling diagram showing EGFR/RAS pathway inhibitors that were assessed for pairwise synergy by isobologram analysis using 50:50 dose-equivalent mixes of each drug pair. **(B)** Heat map of Combination Indices from isobologram analyses of the indicated drug-drug combinations in NT and *SOS2* KO NSCLC cell lines. Synergistic combinations are indicated in yellow, orange, or red. Data are presented as the mean from three independent experiments. **(C-D)** Isobologram analysis and Combination Index (CI) from dose-equivalent treatments of 3D spheroid cultured NCI-H1975 cells treated with the indicated 2-drug **(C)** or 3-drug **(D)** combinations of osimertinib (black), RMC-4550 (purple), and BAY-293 (red). For three drug combination, the two drugs indicated on the y-axis were held at a 1:1 ratio, and then mixed at dose equivalent ratiow with the third drug. CI values indicate enhanced synergy beyond the two drug combination on the y-axis of the isobologram plot and are calculated based on the y-axis drug combination calculated a s single drug treatment. Additive effects occur on the dashed lines of the isobologram plot and have a CI 0.8-1.2 (gray box), whereas synergistic interactions fall below the dashed lines and have a CI < 0.8. **(E)** Combination indices from 2-drug combinations of osimertinib (black), RMC-4550 (purple), and BAY-293 (red) mixed at 2:1, 1:1, or 1:2 ratios or the three drug combination at a 1:1:1 ratio (grey). CI are calculated based on three individual drug treatments. **(F)** Signaling model based on data from Figs. 1–7 showing that combined targeting of mutated EGFR and SOS1 provides sufficient vertical inhibition of upstream signaling to inhibit RAS effector signaling and block oncogenic transformation. This synergistic inhibition can be further enhanced by SHP2 inhibition, providing multiple potential drug combinations for therapeutic intervention in EGFR-mutated NSCLC. Isobologram and CI data are presented as mean +/− s.d. from three independent experiments.

SHP2 is important for the stabilization of the GRB2:SOS1/2 complexes on EGFR (*42*), and the mechanism of allosteric SHP2 inhibitors depends on SOS1 (*43*), although the contribution of SOS2 to SHP2 inhibitors was not assessed. To determine whether *SOS2* deletion altered the spectrum of drug-drug synergies in EGFR-mutated cells, parallel studies were performed in EGFR-mutated cells where *SOS2* was deleted (Fig. 7B, bottom). Unlike what we observed for synergy between EGFR- and SOS1 inhibition, synergy between SOS1 and SHP2 inhibition was enhanced by *SOS2* deletion. These data suggest that SOS2 plays a role in SHP2-dependent signaling. SOS1 inhibition also synergized with MEK inhibition in *SOS2* KO cells. Given the strong synergy between SOS1 inhibition and SOS2 deletion in inhibiting Raf/MEK/ERK signaling (Fig. 6), these data suggest that deep inhibition of MEK signaling is sufficient to inhibit survival in EGFR-mutated cells.

To further evaluate synergy between inhibitors of proximal EGFR signaling, we examined combinations of EGFR-SOS1- and SPH2 inhibition both by expanded evaluation of each 2-drug combination and by assessing whether combined inhibition of EGFR, SOS1, and SHP2 would be more effective than two drug combinations of these inhibitors. To assess each two-drug combination, H1975 cells cultured under 3D spheroid conditions were treated with dose-equivalent combinations of osimertinib, BAY-293, and RMC-4550, assessed for cell viability, and subjected to isobologram analysis to assess drug-drug synergy. Each two-drug combination showed synergy at three different DEQ ratios (Fig. 7C), suggesting that inhibition of any two proximal signaling proteins may be an effective therapeutic regimen to treat EGFR-mutated cancer. To assess whether adding a third proximal inhibitor to each two-drug combination would further enhance synergistic inhibition of spheroid survival, each 2-drug combination was mixed at 1:1 ratio, and then a third proximal pathway inhibitor was added to give the indicated 3-drug mixtures (Fig. 7D). Isobologram analysis of these three drug mixtures revealed that addition of a third proximal pathway inhibitor to any 2-drug combination of osimertinib, BAY-293, and RMC-4550 further enhanced synergy above what was observed for each 2-drug combination (Fig. 7D) Finally, comparing the combination index for the three-drug combination at a 1:1:1 ratio when each drug is treated independently versus the two-drug combinations showed marked synergy for the three drug combination, but that this synergy was not significantly enhanced compared to the combination of osimertinib and BAY-293 (Fig. 7E). These data indicate that vertical inhibition of proximal EGFR signaling with the combination of osimertinib and a SOS1 inhibitor may be the most the most effective therapeutic combination to treat *EGFR*-mutated NSCLC.

## Discussion

Activating *EGFR* mutations are found in 10-15% of lung adenocarcinomas and are the major cause of lung cancer in never smokers. The third-generation EGFR-TKI osimertinib enhances both progression-free (*44*) and overall survival (*45*) compared to first generation EGFR-TKIs and is now considered first-line treatment in EGFR-mutated NSCLC. Osimertinib resistance often develops via activation of parallel RTK pathways (*7–9*), and broad inhibition RTK signaling may enhance osimertinib efficacy and delay therapeutic resistance. Here, we demonstrate that inhibition of the common RTK signaling intermediate SOS1 using BAY-293 showed marked synergy with osimertinib in 3D spheroid-cultured EGFR-mutated NSCLC cells. Our observations that (i) osimertinib–BAY-293 synergy was only observed in 3D spheroids but not in adherent (2D) cultures and (ii) synergy between RTK-signaling intermediates and osimertinib was not broadly applicable EGFR downstream signaling components but was limited to proteins upstream of RAS reveal novel insights into pharmacologic studies assessing therapeutics designed to treat NSCLC.

While most studies designed to identify or test therapeutic targets to treat cancer are done in 2D adherent culture, a growing body of evidence suggests that pharmacologic assessment of novel therapeutics must be performed in 3D culture systems (*34*). This is particularly true for NSCLC, where multiple studies have now revealed the importance of 3D culture systems in order to recapitulate *in vivo* findings. EGFR-mutated cells show differential RTK expression and phosphorylation in 3D versus 2D conditions (*27*) and respond more robustly to EGFR-TKIs in 3D cultures compared to 2D settings (Fig. 1 and (*28*)); KRAS-mutated cell lines deemed “KRAS-independent” in 2D culture (*46–50*) still require KRAS for anchorage-independent growth (*51–54*), and some KRAS^G12C^-mutated NSCLC cell lines respond to KRAS(G12C) inhibitors in 3D culture and *in vivo* but not in 2D adherent culture (*35*). The relevance of 3D culture systems extends to the identification of novel therapeutic targets and therapeutic combinations. We recently showed that SOS2 is specifically required for PI3K-dependent protection from anoikis in KRAS-mutated NSCLC cells (*32*) and *SOS2* deletion synergizes with MEK inhibition to kill *KRAS* mutated cells only under 3D culture conditions (*31*). Here, we show marked synergy between vertical inhibition of EGFR and SOS1 in *EGFR* mutated cancer cells, but only under 3D culture conditions (Fig. 2). CRISPR screens performed in spheroid cultures of KRAS- and EGFR-mutated NSCLC cell lines more accurately reproduce *in vivo* findings and identify drivers of oncogenic growth compared to screens performed in 2D cultures (*55*). Intriguingly, in this study SOS1 was essential for 3D spheroid survival but not 2D spheroid growth of both EGFR- and KRAS-mutated cells. These data are in complete agreement with our data from Fig. 1 showing the requirement for SOS1 in 3D transformation but not 2D proliferation, and support our conclusion that SOS1 is an important therapeutic target in *EGFR*-mutated NSCLC. These data suggest that future studies assessing novel therapeutics to treat lung adenocarcinomas must be performed in a 3D setting.

Osimertinib resistant can occur via oncogenic shift to alternative RTKs including c-MET (*11*), HER2 and/or HER3 (*7–9*), IGF1R (*12*), and AXL (*13–16*). The variety of RTK bypass pathways that can lead to osimertinib resistance suggests that broad inhibition of RTK signaling may be a more effective therapeutic strategy than any individual RTK inhibitor to limit osimertinib resistance, whereas once resistance via oncogenic shift to an alternative RTK occurs then inhibition of the upregulated RTK would have therapeutic benefit. Toward this end, Phase I and II clinical trials are currently examining whether combining osimertinib with inhibitors of AXL (DS-1205c, NCT03255083) or c-MET ((teponitib, NCT03940703; savolitinib, NCT03778229) are effective in patients who have progressed on osimertinib treatment.

Combining osimertinib with a MEK inhibitor can enhance osimertinib efficacy (*10, 17, 20, 56, 57*) and Phase II clinical trials are currently underway to assess combining osimertinib with the MEK inhibitor selumetinib in EGFR-mutated NSCLC (NCT03392246), although resistance to combined osimertinib and MEK inhibition still occurs (*17*). In a recent study designed to understand resistance to combined osimertinib and MEK inhibition, Kurppa et al. (2020) show that combining osimertinib with the MEK inhibitor trametinib results in *EGFR*-mutated cells entering a senescent state that is dependent on the activation of the Hippo pathway effector YAP and its transcription factor binding partner TEAD (*58*). Inhibition of YAP/TEAD signaling overcame this senescence and enhanced killing of EGFR-mutated cells (*58*). EGFR-signaling drives YAP nuclear translocation and transcriptional regulation through PI3K-PDK1 signaling (*59–61*). This suggest that therapeutic combinations able to synergistically inhibit both Raf/MEK/ERK and PI3K/AKT effector signaling should overcome YAP-dependent senescence and treat *EGFR*-mutated NSCLC.

Here, we show that osimertinib does not broadly synergize with inhibitors of downstream EGFR/RAS/RAS effector signaling. Instead, we found that synergy was limited to combinations of osimertinib with inhibitors of proximal EGFR signaling intermediates SOS1 and SHP2 (Fig. 7). Further, SOS1 inhibition significantly enhanced osimertinib-dependent inhibition of both Raf/MEK/ERK and PI3K/AKT signaling (Fig. 6), whereas inhibition of individual downstream Raf/MEK/ERK or PI3K/AKT effector pathways did not synergize with osimertinib (Fig. 7) to inhibit 3D spheroid growth. We hypothesize that these two findings are inexorably linked, so that any potential therapeutic must synergize with osimertinib to inhibit all downstream RAS effector signaling to show drug-drug synergy in 3D culture. In support of this idea, previous studies showed inhibition of SRC family kinases (SFK) potentiated osimertinib to a much greater extent than either MEK or PI3K inhibition (*20*), and that SFK inhibition synergized with osimertinib to inhibit both Raf/MEK/ERK and PI3K/AKT signaling (*20, 62*).

Overall, our data suggest that inhibitors of proximal signaling may be the most efficacious therapeutics to combine with osimertinib to treat EGFR-mutated tumors. Toward this end, Phase I trials are currently underway assessing the combination of osimertinib and the SRC inhibitor dasatinib (NCT02954523) in *EGFR*-mutated NSCLC, and recently developed SOS1 (BI-1701963, NCT04111458) and SHP2 (JAB-3068, NCT03565003; RMC-4630, NCT03634982) inhibitors have entered Phase I safety trials. Our study provides a framework for the systematic, preclinical assessment of therapeutic combinations designed to treat EGFR-mutated cancer cells. We show both how to use basic pharmacologic principles to assess drug-drug synergy and that these combinations must be assessed under 3D culture conditions. Using this framework, we show that the combination of osimertinib and the SOS1 inhibitor BAY-293 shows marked efficacy in 3D spheroid culture and should be pursued as a therapeutic option to treat EGFR-mutated lung adenocarcinoma.

## Acknowledgments

We thank Udayan NCI-H1975, HCC827, PC9, H3255 and H3255-TM cells and for helpful discussions throughout the project. We thank Julian Downward for PC9-TM cells. **Funding**: This work was supported by grants from the Congressionally Directed Medical Research Program to R.L.K. (LC160222 and LC180213). **Author contributions**: P.L.T. and R.L.K. designed the experiments and analyzed the data; P.L.T. and R.L.K. performed most of the experiments; E.S. assisted with isobologram, Bliss independence, and signaling experiments and gave conceptual input throughout the project; A.J.L. performed Western blots and assisted with cell culture and signaling experiments; N.E.S. assisted with isobologram and signaling experiments; D.J.P. assisted with isobologram and transformation experiments. P.L.T. and R.L.K. wrote the manuscript, and E.S. edited the manuscript. **Competing interests**: The authors declare they have no competing interests. **Data and materials availability**: All data needed to evaluate the conclusions in the paper are present in the paper or the Supplementary Materials. Materials are available upon request from R.L.K.

## Materials and Methods

### Cell culture

Cell lines were cultured at 37°C and 5% CO_2_. HCC827, NCI-H1975, PC9, and PC9-TM cells were maintained in Roswell Park Memorial Institute medium (RPMI), each supplemented with 10% fetal bovine serum and 1% penicillin-streptomycin. H3255 and H3255-TM were maintained in ACL4 medium formulated in DMEM:F-12 including: Bovine Serum Albumin 0.5% (w/v) (Sigma cat no. A8022), apo-Transferrin (human) (Sigma cat no. T5391) 0.01 mg/mL, Sodium Selenite (Sigma cat no. S9133) 25nM, Hydrocortisone (Sigma cat no. H0135) 50nM, Ethanolamine (Sigma cat no. E0135) 0.01mM, O-Phosphorylethanolamine (Sigma cat no. P0503) 0.01mM, 3,3’,5-Triiodo-L-thyronine [T3] (Sigma cat no. T5516) 100pM, Sodium Pyruvate (Sigma cat no. P4562), HEPES (Invitrogen cat no 15630-080) 10mM, Epidermal Growth Factor [EGF] 1ng/mL, Recombinant Human Insulin (Sigma cat no. I9278) 0.02mg/mL, and 1% penicillin-streptomycin. For signaling experiments, cells were seeded in 24-well micropatterned AggreWell 400 low-attachment culture plates (Stem Cell # 34415) at 1.2 10^6^ cells/well in 2 mL of medium. 24 h post-plating, half of the media was carefully replaced with fresh media to not disturb the spheroids. At 48 hours, 1 mL media was removed and replaced with 2 x inhibitor. Cells were treated with inhibitor for 6 hrs and then collected for cell lysis and western blot analysis.

### Cell lysis and Western blot analysis

Cells were lysed in RIPA buffer (1% NP-40, 0.1% SDS, 0.1% Na-deoxycholate, 10% glycerol, 0.137 M NaCl, 20 mM Tris pH [8.0], protease (Biotool #B14002) and phosphatase (Biotool #B15002) inhibitor cocktails) for 20 minutes at 4°C and spun at 10,000 RPM for 10 minutes. Clarified lysates were boiled in SDS sample buffer containing 100 mM DTT for 10 minutes prior to Western blotting. Proteins were resolved by sodium dodecyl sulfate-polyacrylamide (Criterion TGX precast) gel electrophoresis and transferred to nitrocellulose membranes. Western blots were developed by multiplex Western blotting using anti-SOS1 (Santa Cruz sc-256; 1:500), anti-SOS2 (Santa Cruz sc-258; 1:500), anti-β-actin (Sigma AC-15; 1:5,000), anti-pEGFR (Cell Signaling 11862; 1:1000), anti-pERK1/2 (Cell Signaling 4370; 1:1,000), anti-ERK1/2 (Cell Signaling 4696; 1:1000), anti-pAKT Ser^473^ [Cell Signaling 4060; 1:1000]), anti-AKT (Cell Signaling 2920; 1:1000), anti-HSP90 (Santa Crux sc-7947, 1:1000), primary antibodies. Anti-mouse and anti-rabbit secondary antibodies conjugated to IRDye680 or IRDye800 (LI-COR; 1:10,000) were used to probe primary antibodies. Western blot protein bands were detected and quantified using the Odyssey system (LI-COR). For quantification of SOS1 and SOS2 abundance, samples were normalized to either β-actin or HSP90. For quantification of pERK and pAKT, samples were normalized to a weighted average of HSP90, β-actin, total ERK1/2, total AKT, and total EGFR (*63*).

### Proliferation Studies

For 2D proliferationh assays, 5 × 10^2^ cells were seeded on cell culture-coated 96-well white-walled CulturePlates (Perkin Elmer #6005688). Cells were lysed with CellTiter-Glo® 2.0 Reagent (Promega), and luminescence was read using a Bio-Tek Cytation 5 multi-mode plate reader. Cell number was assessed 24 hours after plating to account for any discrepancies in plating (Day 1), and then on days 3, 5, and 7. Data were analyzed as an increase in luminescence over Day 1.

### Transformation Studies

H3255 and H3255-TM cells were seeded in 0.32% Nobel agar at 2 × 10^4^ cells per 35-mm dish to assess anchorage-independent. Soft agar colonies were counted 28 days after seeding. For all other cell lines spheroid growth assessed in ultra-low attachment 96-well round bottomed plates (Corning Costar #7007), cells were seeded at 500 cells per well. Images were taken 24 hours after plating to assess initial spheroid size, and then 7, 14, and 21 days later to assess transformation. Cell number was assessed in parallel plates at 0, 7, 14, and 21 days using CellTiter-Glo® 2.0 reagent.

### sgRNA studies

A non-targeting (NT) single guide RNA (sgRNA), a SOS2-targeted sgRNA (*31*), and 8 potential SOS1-targeted sgRNAs previously used to target SOS1 in a genome-wide CRISPR screen (*37*) were each cloned into pLentiCRISPRv2 as previously described (*64*). SOS1-2 was chosen as the SOS1 sgRNA for the study.

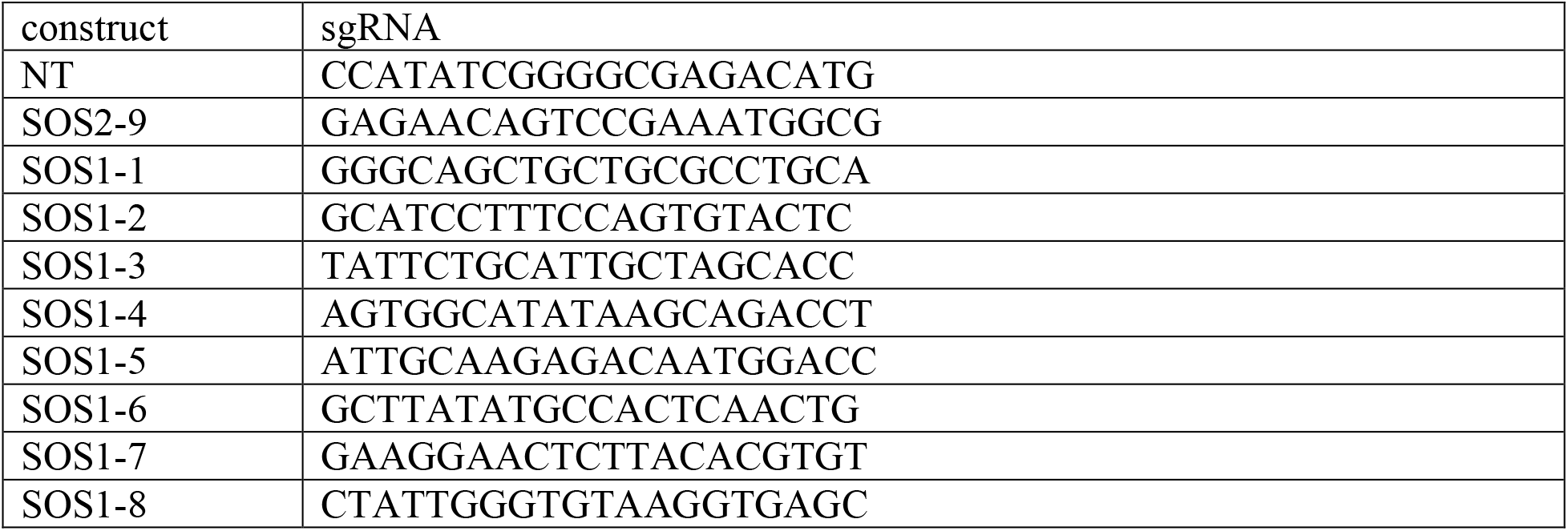

### Production of recombinant lentiviruses

Lentiviruses were produced by co-transfecting MISSION lentiviral packaging mix (Sigma) into 293T cells using Mirus *Trans*IT^®^-Lenti transfection reagent (Mirus Bio # MIR6605) in Opti-MEM (Thermo Scientific #31-985-062). At 48 h post-transfection, viral supernatants were collected and filtered. Viral supernatants were then either stored at −80°C or used immediately to infect cells in combination with polybrene at 8 μg/mL. 48 hours post-infection, cells were selected in 4 μg/mL Puromycin (Invitrogen). Twelve days after selection, cells were analyzed for SOS1 and SOS2 expression and plated for proliferation and transformation assays.

### Inhibitor Studies

- 2D adherent studies – Cells were seeded at 500-1,000 cells per well in 100 μL in the inner-60 wells of 96-well white-walled culture plates (Perkin Elmer) and allowed to attach for 48 hours prior to drug treatment. Cells were treated with drug for 72 hours prior to assessment of cell viability using CellTiter-Glo® 2.0.
- 3D adherent studies – Cells were seeded at 500-1,000 cells per well in 100 μL in the inner-60 wells of 96-well ultra-low attachment round bottomed plates (Corning #7007) or Nunc Nucleon Sphera microplates (ThermoFisher # 174929) and allowed to coalesce as spheroids for 48-72 hours prior to drug treatment. Cells were treated with drug for 96 hours prior to assessment of cell viability using CellTiter-Glo® 2.0.

For all studies, outer wells (rows A and H, columns 1 and 12) were filled with 200 μL of PBS to buffer inner cells from temperature and humidity fluctuations. Triplicate wells of cells were then treated with increasing concentrations 100 μL of 2× inhibitor at either a semilog (single drug dose response curves to determine EC_50_) or a 1/3-log scale (isobologram and Bliss independence experiments) for 72 (adherent cultures) or 96 (spheroids) hours. Cell viability was assessed using CellTiter-Glo® 2.0 (30 μL/well). Luminescence was assessed using a Bio-Tek Cytation 5 multi-mode plate reader. Data were normalized to the maximum luminescence reading of untreated cells, and individual drug EC_50_ values were calculated using Prism 8 by non-linear regression using log(inhibitor) vs. response with a variable slope (four parameters) to assess for differences in the Hill Coefficient between different drug treatments.

### Isobologram Analysis

Dose equivalence was first determined by assessing individual-drug EC_50_ values; individual-drug Hill Coefficients were determined to assure that the two drugs could be assessed for synergy by Lowe additivity. To generate dose-equivalent dose-response curves, the dose for each drug closest to the EC_50_ on a 1/3-log scale was set as equivalent, and 10-point dose response curves were generated for each individual drug on either side of the equivalent dose to ensure the top (no drug effect) and bottom (maximal drug effect) were represented on the dose-response curve. 100 μL of drug each drug dose was added as outlined above. To generate dose-equivalent mixtures for isobologram analysis, equivalent doses of the two drugs were mixed at different ratios so that the total dose (100 μL) would be expected to have an equivalent effect on the cells if the two drugs were additive. Drugs were mixed at either five (4:1, 2:1, 1:1, 1:2, and 1:4) or three (2:1, 1:1, and 1:2) different drug mixtures depending on the experiment. Cells were treated and EC_50_ values for each individual drug or drug mixture based on each drug’s dosing were determined for as outlined above. To generate an isobologram plot, the EC_50_ of each individual drug was plotted as the x- or y-intercept, and the calculated contribution of each drug to the overall EC_50_ for each DEQ mix is plotted as a single point (EC_50,A_, EC_50,B_) on the graph.

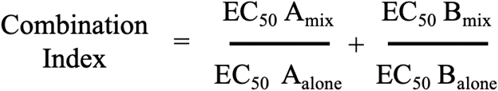

To calculate the combination index for each dose equivalent mixture, the calculated contribution of each drug to the overall EC_50_ were used in the equation:

As an example, we will show data for one trial analyzing the combination of osimertinib and BAY-293 in 3D spheroid cultured H1975 cells in Fig. 2B. The EC_50_ values for each individual drug were first determined: −8.57 for osimertinib and −5.73 for BAY-293. Based on these EC_50_ values, the dose equivalence was set at −5.67 for osimertinib −5.67 for BAY-293 (**approximated EC_50_ for each drug in bold**), and the following 10-point dose response curves were generated:

|  |  |  |  |  |  |  |  |  |  |  |
| --- | --- | --- | --- | --- | --- | --- | --- | --- | --- | --- |
| osimertinib | -11 | -10.67 | -10.33 | -10 | -9.67 | -9.33 | -9 | **-8.67** | -8.33 | -8 |
| BAY-293 | -8 | -7.67 | -7.33 | -7 | -6.67 | -6.33 | -6 | **-5.67** | -5.33 | -5 |

Cells were then treated with the following volumes of each drug to generate seven dose-equivalent dose response curves:

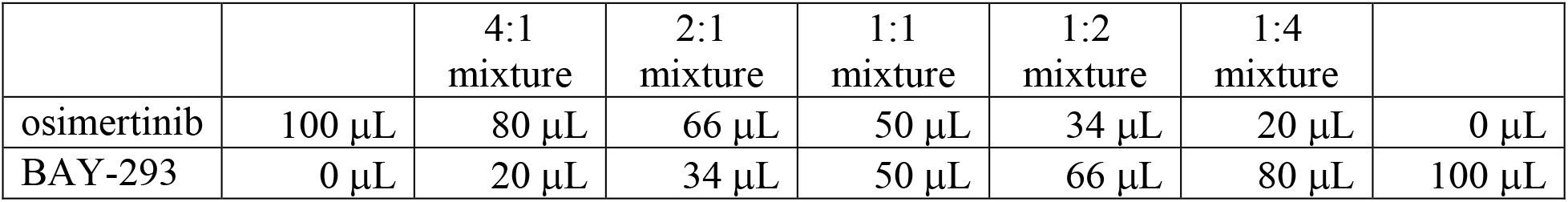

EC_50_ values for each dose-response curve were then determined based on each drug’s dosing:

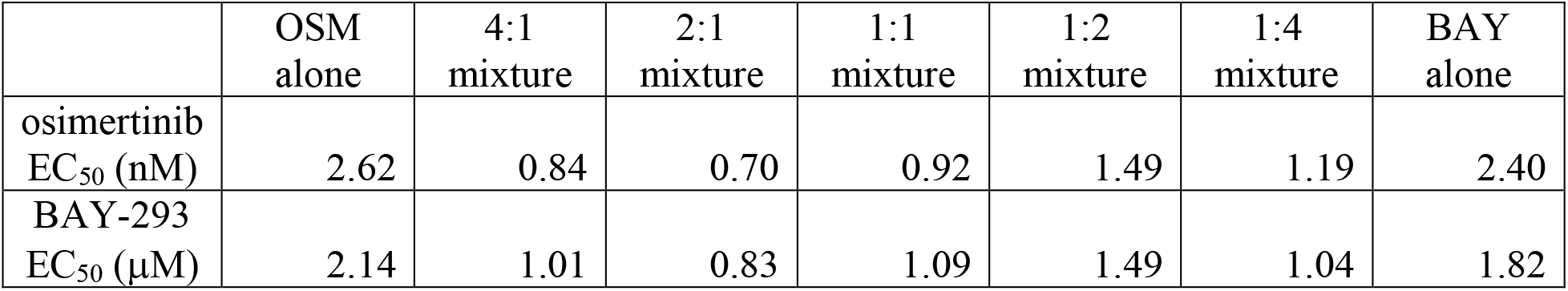

EC_50_ values were then adjusted based on the amount of each drug that was put in the mixture to determine the contribution of each drug in the mixture to the overall EC_50_. For example, the 4:1 mixture was 80% osimertinib, so the osimertinib EC50 for that mixture is multiplied by 0.8. The corresponding corrected EC_50_ values and combination indices were:

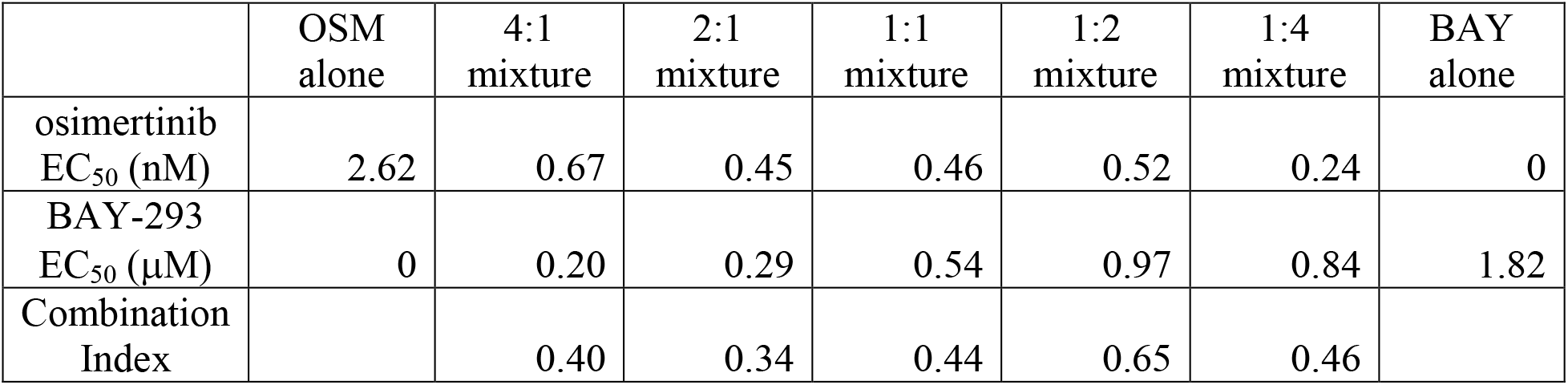

### Bliss Independence Analysis

Unlike Isobologram analysis, individual drug doses are not reduced for drug-drug combinations when performing Bliss independence analysis. For data in Fig. 2, wells were treated with a full dose of each individual drug or drug combination in a 10 × 10 matrix of dose combinations for osimertinib and BAY-293 on a 1/3-log scale. Data were normalized to the maximum luminescence reading of untreated cells, and a heat-map depicting cell viability was generated using Prism 8. The Bliss index was calculated by first converting viability (on a scale of 0 to 1) for each treatment to the effect of each drug or drug combination, where 0 represents no effect and 1 represents 100% effect (no viable cells).

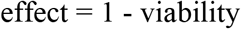

From the effect data, the expected effect for each drug combination is calculated:

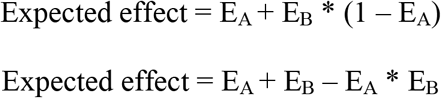

The <u>Bliss Index</u> is the ratio of the expected effect / actual effect:

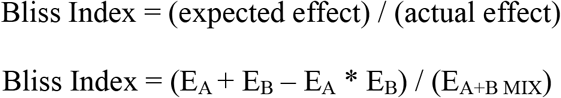

A Bliss Index of 1 indicates that the actual and expected effects are equivalent, and the effects of the two drugs are additive. Bliss Index < 1 indicates increasing synergy, whereas Bliss Index >1 indicates antagonism.

<u>Excess over Bliss</u> is calculated by determining how much greater the actual effect of the drug combination is versus the expected effect, and is calculated as:

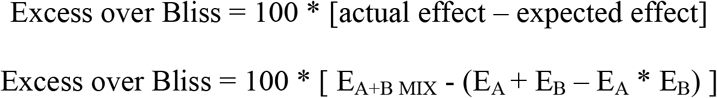

An excess over Bliss of 0 indicates that the actual and expected effects are equivalent, and the effects of the two drugs are additive; values > 0 indicate increasing synergy, whereas values < 0 indicate antagonism.

Since synergy occurred at drug combinations at or just below the EC_50_ values for each individual drug, Bliss experiments in Figs. 4 and 5, drug mixtures were limited to 3 × 10 drug mixtures based on dose equivalence with mixtures at approximately 2:1, 1:1, and 1:2 mixes of the two drugs based on dose equivalence. Here, the doses used for one drug were held constant, and the second drug dose wash shifted by 1/3 log up or down to generate 2:1 and 1:2 mixtures. For example, for the combination of osimertinib and BAY-293 in H1975 cells, the following drug doses were used:

|  |  |  |  |  |  |  |  |  |  |  |
| --- | --- | --- | --- | --- | --- | --- | --- | --- | --- | --- |
| **Osimertinib**<br>(1:2 ratio of<br>OSM:BAY) | -11.33 | -11 | -10.67 | -10.33 | -10 | -9.67 | -9.33 | -9 | -8.67 | -8.33 |
| **Osimertinib**<br>(1:1 ratio of<br>OSM:BAY) | -11 | -10.67 | -10.33 | -10 | -9.67 | -9.33 | -9 | -8.67 | -8.33 | -8 |
| **Osimertinib**<br>(2:1 ratio of<br>OSM:BAY) | -10.67 | -10.33 | -10 | -9.67 | -9.33 | -9 | -8.67 | -8.33 | -8 | -7.67 |
| **BAY-293**<br>(constant) | -8 | -7.67 | -7.33 | -7 | -6.67 | -6.33 | -6 | -5.67 | -5.33 | -5 |

### 3-drug Isobologram analysis

For three-drug isobologram studies with osimertinib (EC_50_ = −5.57), BAY-293 (EC_50_ = −5.74), and RCM-4550 (EC_50_ = −5.84), drugs were again mixed based on dose equivalency. The dose-equivalent 10-point dose-response curves for these drugs in 3D cultured H1975 cells were (**approximated EC_50_ for each drug in bold**):

|  |  |  |  |  |  |  |  |  |  |  |
| --- | --- | --- | --- | --- | --- | --- | --- | --- | --- | --- |
| osimertinib | -11 | <sup>-</sup><br>10.67 | <sup>-</sup><br>10.33 | -10 | -9.67 | -9.33 | -9 | **-8.67** | -8.33 | -8 |
| BAY-293 | -8 | -7.67 | -7.33 | -7 | -6.67 | -6.33 | -6 | **-5.67** | -5.33 | -5 |
| RCM-4550 | -9 | **-8.67** | -8.33 | -8 | -7.67 | -7.33 | -7 | **-6.67** | -6.33 | -6 |

Each two-drug combination was set as a single “drug mixture” at a 1:1 ratio, and the third drug was combined with this drug mixture at 2:1, 1:1, and 1:2 drug ratios. To generate the proper two and three-drug mixtures for analysis, 21 total dose response curves were generated. The five dose-response curves on the right represent the mixtures used to generate the isobologram plots in Fig. 7D. The other two two-drug mixtures in **bold** (2-drug 2:1 and 1:2 mixtures) were used to generate the isobologram plots in Fig. 7CCombination indices were calculated based on whether addition of the third drug to each 2-drug 1:1 mixture further enhanced synergy when added to the two-drug mixture.

[osimertinib:BAY-293] mixture vs. RCM-4550.

[osimertinib:BAY-293] mixture vs. RCM-4550
|  | **OSM:BAY**<br>**2:1** | **OSM:BAY**<br>**1:2** | OSM:BAY<br>1:1 | (1+1):1<br>2:1<br>mixture | (1+1):2<br>1:1<br>mixture | (1+1):4<br>1:2<br>mixture | RCM<br>alone |
| --- | --- | --- | --- | --- | --- | --- | --- |
| osimertinib | **66** μL | **34** μL | 50 μL | 33 μL | 25 μL | 17 μL | 0 μL |
| BAY-293 | **34** μL | **66** μL | 50 μL | 33 μL | 25 μL | 17 μL | 0 μL |
| RCM-4550 | **0** μL | **0** μL | 0 μL | 34 μL | 50 μL | 66 μL | 100 μL |

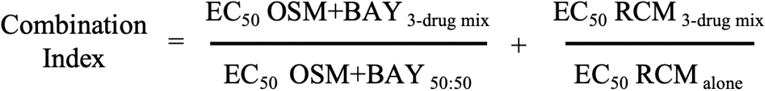

[osimertinib:RCM-4550] mixture vs. BAY-293.

|  | **OSM:RCM**<br>**2:1** | **OSM:RCM**<br>**1:2** | **OSM:RCM**<br>**1:1** | **(1+1):1**<br>**2:1**<br>**mixture** | **(1+1):2**<br>**1:1**<br>**mixture** | **(1+1):4**<br>**1:2**<br>**mixture** | **RCM**<br>**alone** |
| --- | --- | --- | --- | --- | --- | --- | --- |
| osimertinib | **66 µL** | **34 µL** | 50 µL | 33 µL | 25 µL | 17 µL | 0 µL |
| BAY-293 | **0 µL** | **0 µL** | 0 µL | 34 µL | 50 µL | 66 µL | 100 µL |
| RMC-4550 | **34 µL** | **66 µL** | 50 µL | 33 µL | 25 µL | 17 µL | 0 µL |

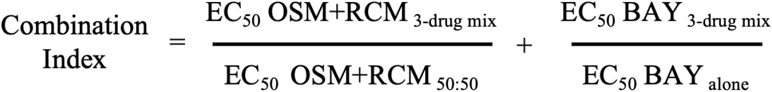

[BAY-293:RCM-4550] mixture vs. osimertinib.

|  | **BAY:RCM**<br>**2:1** | **BAY:RCM**<br>**1:2** | **BAY:RCM**<br>**1:1** | **(1+1):1**<br>**2:1**<br>**mixture** | **(1+1):2**<br>**1:1**<br>**mixture** | **(1+1):4**<br>**1:2**<br>**mixture** | **RCM**<br>**alone** |
| --- | --- | --- | --- | --- | --- | --- | --- |
| osimertinib | **0 µL** | **0 µL** | 0 µL | 34 µL | 50 µL | 66 µL | 100 µL |
| BAY-293 | **66 µL** | **34 µL** | 50 µL | 33 µL | 25 µL | 17 µL | 0 µL |
| RMC-4550 | **34 µL** | **66 µL** | 50 µL | 33 µL | 25 µL | 17 µL | 0 µL |

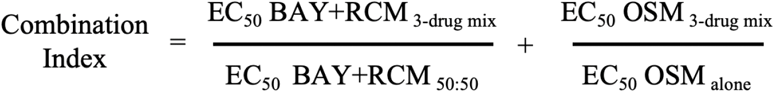

To calculate the three-drug combination index where each drug was considered independently (Fig. 7E), the following equation was used:

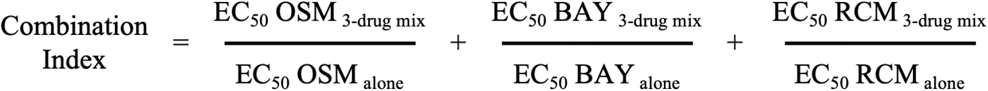

## References

1. N. Cancer Genome Atlas Research, Comprehensive molecular profiling of lung adenocarcinoma. Nature 511, 543–550 (2014).

2. T. S. Mok et al., Gefitinib or carboplatin-paclitaxel in pulmonary adenocarcinoma. N Engl J Med 361, 947–957 (2009).

3. J. J. Yang et al., A phase III randomised controlled trial of erlotinib vs gefitinib in advanced non-small cell lung cancer with EGFR mutations. Br J Cancer 116, 568–574 (2017).

4. D. A. Eberhard et al., Mutations in the epidermal growth factor receptor and in KRAS are predictive and prognostic indicators in patients with non-small-cell lung cancer treated with chemotherapy alone and in combination with erlotinib. J Clin Oncol 23, 5900–5909 (2005).

5. P. A. Janne et al., AZD9291 in EGFR inhibitor-resistant non-small-cell lung cancer. N Engl J Med 372, 1689–1699 (2015).

6. D. A. Cross et al., AZD9291, an irreversible EGFR TKI, overcomes T790M-mediated resistance to EGFR inhibitors in lung cancer. Cancer Discov 4, 1046–1061 (2014).

7. M. Mancini et al., An oligoclonal antibody durably overcomes resistance of lung cancer to third-generation EGFR inhibitors. EMBO Mol Med 10, 294–308 (2018).

8. D. Romaniello et al., A Combination of Approved Antibodies Overcomes Resistance of Lung Cancer to Osimertinib by Blocking Bypass Pathways. Clin Cancer Res 24, 5610–5621 (2018).

9. S. La Monica et al., Trastuzumab emtansine delays and overcomes resistance to the third-generation EGFR-TKI osimertinib in NSCLC EGFR mutated cell lines. J Exp Clin Cancer Res 36, 174 (2017).

10. C. A. Eberlein et al., Acquired Resistance to the Mutant-Selective EGFR Inhibitor AZD9291 Is Associated with Increased Dependence on RAS Signaling in Preclinical Models. Cancer Res 75, 2489–2500 (2015).

11. p. Shi et al., Met gene amplification and protein hyperactivation is a mechanism of resistance to both first and third generation EGFR inhibitors in lung cancer treatment. Cancer Lett 380, 494–504 (2016).

12. J. H. Park et al., Activation of the IGF1R pathway potentially mediates acquired resistance to mutant-selective 3rd-generation EGF receptor tyrosine kinase inhibitors in advanced non-small cell lung cancer. Oncotarget 7, 22005–22015 (2016).

13. D. Kim et al., AXL degradation in combination with EGFR-TKI can delay and overcome acquired resistance in human non-small cell lung cancer cells. Cell Death Dis 10, 361 (2019).

14. H. Taniguchi et al., AXL confers intrinsic resistance to osimertinib and advances the emergence of tolerant cells. Nat Commun 10, 259 (2019).

15. T. Jimbo et al., DS-1205b, a novel selective inhibitor of AXL kinase, blocks resistance to EGFR-tyrosine kinase inhibitors in a non-small cell lung cancer xenograft model. Oncotarget 10, 5152–5167 (2019).

16. K. Namba et al., Activation of AXL as a Preclinical Acquired Resistance Mechanism Against Osimertinib Treatment in EGFR-Mutant Non-Small Cell Lung Cancer Cells. Mol Cancer Res 17, 499–507 (2019).

17. E. M. Tricker et al., Combined EGFR/MEK Inhibition Prevents the Emergence of Resistance in EGFR-Mutant Lung Cancer. Cancer Discov 5, 960–971 (2015).

18. K. Jacobsen et al., Convergent Akt activation drives acquired EGFR inhibitor resistance in lung cancer. Nat Commun 8, 410 (2017).

19. B. M. Ku et al., Acquired resistance to AZD9291 as an upfront treatment is dependent on ERK signaling in a preclinical model. PLoS One 13, e0194730 (2018).

20. E. Ichihara et al., SFK/FAK Signaling Attenuates Osimertinib Efficacy in Both Drug-Sensitive and Drug-Resistant Models of EGFR-Mutant Lung Cancer. Cancer Res 77, 2990–3000 (2017).

21. A. S. Nunes, A. S. Barros, E. C. Costa, A. F. Moreira, I. J. Correia, 3D tumor spheroids as in vitro models to mimic in vivo human solid tumors resistance to therapeutic drugs. Biotechnol Bioeng 116, 206–226 (2019).

22. S. A. Langhans, Three-Dimensional in Vitro Cell Culture Models in Drug Discovery and Drug Repositioning. Front Pharmacol 9, 6 (2018).

23. H. X. Hao et al., Tumor Intrinsic Efficacy by SHP2 and RTK Inhibitors in KRAS-Mutant Cancers. Mol Cancer Ther 18, 2368–2380 (2019).

24. J. W. Kim, W. J. Ho, B. M. Wu, The role of the 3D environment in hypoxia-induced drug and apoptosis resistance. Anticancer Res 31, 3237–3245 (2011).

25. A. Riedl et al., Comparison of cancer cells in 2D vs 3D culture reveals differences in AKT-mTOR-S6K signaling and drug responses. J Cell Sci 130, 203–218 (2017).

26. G. G. Jones et al., SHOC2 phosphatase-dependent RAF dimerization mediates resistance to MEK inhibition in RAS-mutant cancers. Nat Commun 10, 2532 (2019).

27. J. E. Ekert et al., Three-dimensional lung tumor microenvironment modulates therapeutic compound responsiveness in vitro--implication for drug development. PLoS One 9, e92248 (2014).

28. N. Jacobi et al., Organotypic three-dimensional cancer cell cultures mirror drug responses in vivo: lessons learned from the inhibition of EGFR signaling. Oncotarget 8, 107423–107440 (2017).

29. D. Vigil, J. Cherfils, K. L. Rossman, C. J. Der, Ras superfamily GEFs and GAPs: validated and tractable targets for cancer therapy? Nat Rev Cancer 10, 842–857 (2010).

30. H. H. Jeng, L. J. Taylor, D. Bar-Sagi, Sos-mediated cross-activation of wild-type Ras by oncogenic Ras is essential for tumorigenesis. Nat Commun 3, 1168 (2012).

31. E. Sheffels et al., Oncogenic RAS isoforms show a hierarchical requirement for the guanine nucleotide exchange factor SOS2 to mediate cell transformation. Sci Signal 11, (2018).

32. E. Sheffels, N. E. Sealover, P. L. Theard, R. L. Kortum, Anchorage-independent growth conditions reveal a differential SOS2 dependence for transformation and survival in RAS-mutant cancer cells. Small GTPases, 1–12 (2019).

33. R. C. Hillig et al., Discovery of potent SOS1 inhibitors that block RAS activation via disruption of the RAS-SOS1 interaction. Proc Natl Acad Sci U S A 116, 2551–2560 (2019).

34. A. S. Nunes, A. S. Barros, E. C. Costa, A. F. Moreira, I. J. Correia, 3D tumor spheroids as in vitro models to mimic in vivo human solid tumors resistance to therapeutic drugs. Biotechnology and Bioengineering 116, 206–226 (2019).

35. M. R. Janes et al., Targeting KRAS Mutant Cancers with a Covalent G12C-Specific Inhibitor. Cell 172, 578–589 e517 (2018).

36. N. Jacobi et al., Organotypic three-dimensional cancer cell cultures mirror drug responses in vivo: lessons learned from the inhibition of EGFR signaling. Oncotarget 8, 107423–107440 (2017).

37. D. M. Munoz et al., CRISPR Screens Provide a Comprehensive Assessment of Cancer Vulnerabilities but Generate False-Positive Hits for Highly Amplified Genomic Regions. Cancer Discov 6, 900–913 (2016).

38. E. C. de Bruin et al., Reduced NF1 expression confers resistance to EGFR inhibition in lung cancer. Cancer Discov 4, 606–619 (2014).

39. J. A. Engelman et al., Allelic dilution obscures detection of a biologically significant resistance mutation in EGFR-amplified lung cancer. J Clin Invest 116, 2695–2706 (2006).

40. K. R. Roell, D. M. Reif, A. A. Motsinger-Reif, An Introduction to Terminology and Methodology of Chemical Synergy-Perspectives from Across Disciplines. Front Pharmacol 8, 158 (2017).

41. R. J. Tallarida, Quantitative methods for assessing drug synergism. Genes Cancer 2, 1003–1008 (2011).

42. M. Dance, A. Montagner, J.-p. Salles, A. Yart, p. Raynal, The molecular functions of Shp2 in the Ras/Mitogen-activated protein kinase (ERK1/2) pathway. Cell Signal 20, 453–459 (2008).

43. R. J. Nichols et al., RAS nucleotide cycling underlies the SHP2 phosphatase dependence of mutant BRAF-, NF1- and RAS-driven cancers. Nat Cell Biol 20, 1064–1073 (2018).

44. J. C. Soria et al., Osimertinib in Untreated EGFR-Mutated Advanced Non-Small-Cell Lung Cancer. N Engl J Med 378, 113–125 (2018).

45. S. S. Ramalingam et al., Overall Survival with Osimertinib in Untreated, EGFR-Mutated Advanced NSCLC. N Engl J Med 382, 41–50 (2020).

46. O. A. Balbin et al., Reconstructing targetable pathways in lung cancer by integrating diverse omics data. Nat Commun 4, 2617 (2013).

47. A. Singh et al., A gene expression signature associated with “K-Ras addiction” reveals regulators of EMT and tumor cell survival. Cancer Cell 15, 489–500 (2009).

48. A. Singh et al., TAK1 inhibition promotes apoptosis in KRAS-dependent colon cancers. Cell 148, 639–650 (2012).

49. C. Scholl et al., Synthetic lethal interaction between oncogenic KRAS dependency and STK33 suppression in human cancer cells. Cell 137, 821–834 (2009).

50. S. Lamba et al., RAF suppression synergizes with MEK inhibition in KRAS mutant cancer cells. Cell Rep 8, 1475–1483 (2014).

51. S. Fujita-Sato et al., Enhanced MET Translation and Signaling Sustains K-Ras-Driven Proliferation under Anchorage-Independent Growth Conditions. Cancer Res 75, 2851–2862 (2015).

52. A. Rotem et al., Alternative to the soft-agar assay that permits high-throughput drug and genetic screens for cellular transformation. Proc Natl Acad Sci U S A 112, 5708–5713 (2015).

53. Z. Zhang, G. Jiang, F. Yang, J. Wang, Knockdown of mutant K-ras expression by adenovirus-mediated siRNA inhibits the in vitro and in vivo growth of lung cancer cells. Cancer Biol Ther 5, 1481–1486 (2006).

54. F. McCormick, KRAS as a Therapeutic Target. Clin Cancer Res 21, 1797–1801 (2015).

55. K. Han et al., CRISPR screens in cancer spheroids identify 3D growth-specific vulnerabilities. Nature ePub March 11, 2020, (2020).

56. p. Shi et al., Overcoming Acquired Resistance to AZD9291, A Third-Generation EGFR Inhibitor, through Modulation of MEK/ERK-Dependent Bim and Mcl-1 Degradation. Clin Cancer Res 23, 6567–6579 (2017).

57. C. M. Della Corte et al., Antitumor Efficacy of Dual Blockade of EGFR Signaling by Osimertinib in Combination With Selumetinib or Cetuximab in Activated EGFR Human NCLC Tumor Models. J Thorac Oncol 13, 810–820 (2018).

58. K. J. Kurppa et al., Treatment-Induced Tumor Dormancy through YAP-Mediated Transcriptional Reprogramming of the Apoptotic Pathway. Cancer Cell 37, 104–122 e112 (2020).

59. R. Fan, N. G. Kim, B. M. Gumbiner, Regulation of Hippo pathway by mitogenic growth factors via phosphoinositide 3-kinase and phosphoinositide-dependent kinase-1. Proc Natl Acad Sci U S A 110, 2569–2574 (2013).

60. H. Xia et al., EGFR-PI3K-PDK1 pathway regulates YAP signaling in hepatocellular carcinoma: the mechanism and its implications in targeted therapy. Cell Death Dis 9, 269 (2018).

61. K. Tumaneng et al., YAP mediates crosstalk between the Hippo and PI(3)K-TOR pathways by suppressing PTEN via miR-29. Nat Cell Biol 14, 1322–1329 (2012).

62. S. Watanabe et al., T790M-Selective EGFR-TKI Combined with Dasatinib as an Optimal Strategy for Overcoming EGFR-TKI Resistance in T790M-Positive Non-Small Cell Lung Cancer. Mol Cancer Ther 16, 2563–2571 (2017).

63. K. A. Janes, An analysis of critical factors for quantitative immunoblotting. Sci Signal 8, rs2 (2015).

64. N. E. Sanjana, O. Shalem, F. Zhang, Improved vectors and genome-wide libraries for CRISPR screening. Nat Methods 11, 783–784 (2014).

