## Supplemental Figures 1-3 for "Marked Synergy by Vertical Inhibition of EGFR signaling in NSCLC Spheroids: SOS1 as a therapeutic target in EGFR-mutated cancer"

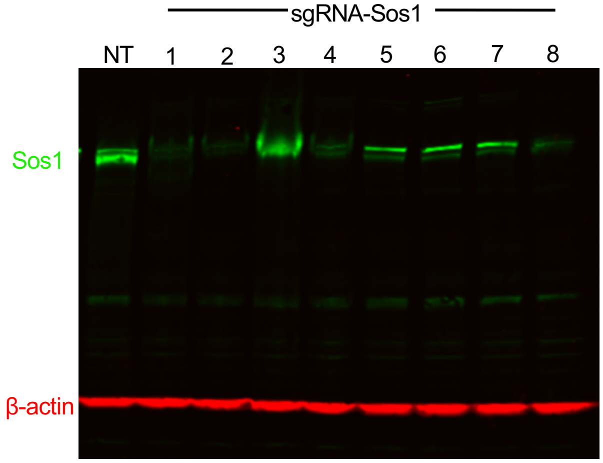


Figure S1. **Deletion of SOS1 using CRISPR/Cas9.**
293T cells were transduced with lentiviruses expressing Cas9 and either a non-targeting sgRNA (NT) or one of eight different sgRNAs targeting SOS1. Whole cell lysates (WCLs) were analyzed by Western blotting with antibodies specific for SOS1 or β-actin. SOS1 sgRNA constructs #1, #2, and #8 consistently showed >90% reduction in total SOS1 protein abundance. SOS1-2 was used for experiments in Fig. 1.

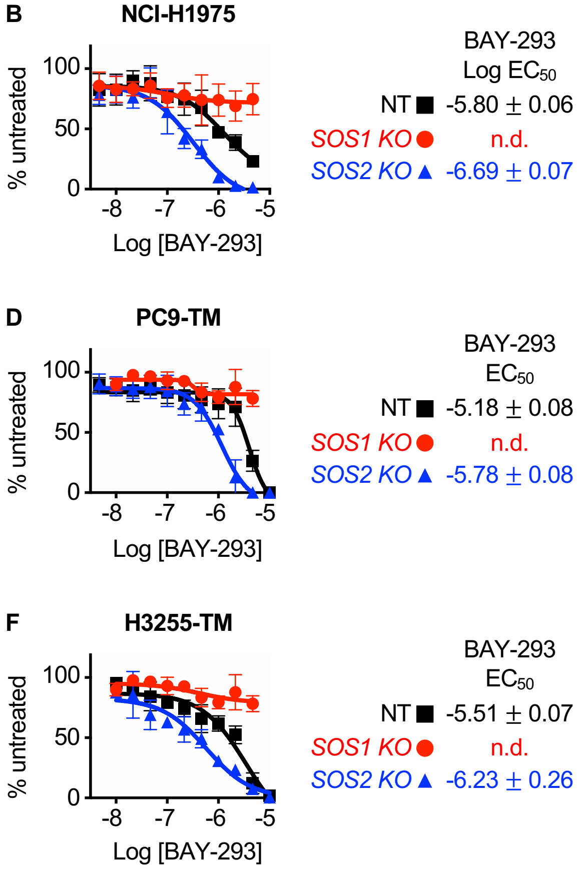


Figure S2. **The SOS1 inhibitor BAY-293 is specific for SOS1 and is enhanced by *SOS2* deletion in EGFR (T790M) mutated NSCLC cell lines.**
A-C. Dose-response curves of NCI-H1975 (A), PC9-TM (B), or H3255-TM (C) cells where *SOS1* (red circles) or *SOS2* (blue triangles) has been deleted using CRISPR/Cas9 vs NT controls (black squares) treated with BAY-293 under 3D spheroid culture conditions.
Data are presented as mean +/- s.d. from at least three independent experiments.

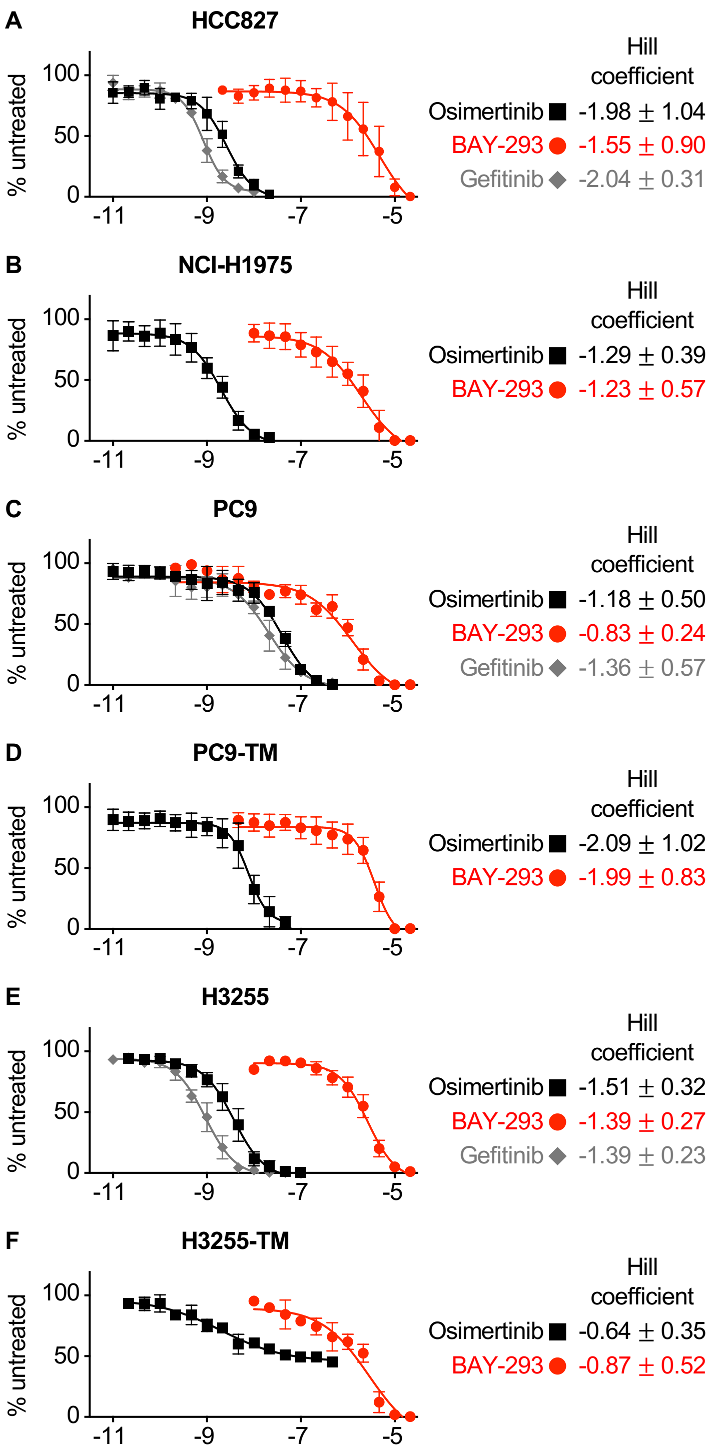


Figure S3. **EGFR mutated NSCLC cell lines are responsive to osimertinib, BAY-293, and gefitinib in 3D spheroid cultures.**
A-F. Dose-response curves of 3D spheroid cultured HCC827 (A), NCI-H1975 (B), PC9 (C), PC9-TM (D), H3255 (E), or H3255-TM (F) cells to osimertinib (black squares), BAY-293 (red circles) or gefitinib (grey diamonds). Hill coefficients +/- s.d. are shown to the right.
Data are presented as mean +/- s.d. from at least three independent experiments.
